# Rapid enzymatic detection of Shigatoxin-producing *E. coli* using fluorescence-labeled oligonucleotide substrates

**DOI:** 10.1101/2023.10.12.562006

**Authors:** Isabell Ramming, Christina Lang, Samuel Hauf, Maren Krüger, Sylvia Worbs, Carsten Peukert, Angelika Fruth, Brigitte G. Dorner, Mark Brönstrup, Antje Flieger

## Abstract

Shigatoxin-producing *E. coli* (STEC) are important human pathogens causing disease ranging from diarrhea to severe hemolytic uremic syndrome. As STEC are transmitted via animals, food, and water, and may produce large outbreaks, their timely and qualified detection including isolate recovery is of high importance, but challenging and labor-intense. Thus, the availability of an easy-to-perform rapid test would be a tremendous advance. Since the common feature and major virulence factor of otherwise multifaceted STEC is the Shiga toxin (Stx), we developed a detection method for Stx, specifically for its catalytic RNA-*N*-glycosidase activity targeting the Sarcin Ricin Loop (SRL) of 28S ribosomal RNA. To this end, synthetic ssDNA substrates mimicking the SRL were designed and linked to a fluorophore and quencher pair, which conferred a fluorescence signal after cleavage by Stx. Optimal results using bacterial culture supernatants or single colonies were achieved for substrate **StxSense 4** following 30 to 60 minutes incubation. Importantly, different Stx1 and Stx2 subtypes, diverse STEC serotypes, and *Shigella* were detected. In conclusion, the assay offers rapid and facile detection of STEC based on a real-time readout for Stx activity. Therefore, it may improve STEC risk evaluation, therapy decisions, outbreak and source detection, and simplify research for antimicrobials.

## Introduction

The species *Escherichia coli* is on the one hand part of the commensal intestinal microbiome, and comprises on the other hand pathovars causing disease, such as Shiga toxin-producing *E. coli* (STEC). STEC may cause a range of symptoms including diarrhea, bloody diarrhea, and the severe hemolytic uremic syndrome (HUS) affecting predominantly children up to the age of five. STEC are important zoonotic pathogens and are found in association with animals, such as ruminants, and food, especially meat and milk, but also plant-derived products ^1^. They cause high socio-economic and economic costs due to their ability to generate large outbreaks, such as the STEC O104:H4 outbreak in 2011 encompassing ∼3000 cases of diarrhea, more than 800 cases of HUS, and 54 fatalities ^2^.

STEC possess a variety of virulence factors but Shiga toxin (Stx), an AB5 toxin showing RNA-*N-* glycosidase activity, is the most significant virulence factor and is found in all of the diverse STEC ^3^. STEC Stx divides into two major groups, Stx1 and Stx2, which share ∼50 % protein sequence identity (Tab. S1). Stx1 is more closely related to Stx of *S. dysenteriae* than Stx2. So far, Stx subtypes Stx1a, Stx1c, and Stx1d and Stx2a-o are known in STEC (Tab. S1) ^4,5^. Stx2 and the subtypes Stx2a, Stx2c, and Stx2d are most frequently associated with HUS and severe disease ^6^.

In AB5 toxins, the five Stx B subunits bind to specific receptors and the A subunit confers the enzymatic activity. The action of Stx in the eukaryotic cell is based on the binding of the B subunit to cellular ganglioside Gb3/CD77 receptors as found on renal cells, subsequent endocytosis, and release of the furin-processed active enzyme A1 subunit into the host cell (Fig. S1) ^3^. Subunit A1 cleaves a single adenine from the 28S ribosomal RNA (rRNA) located in the Sarcin Ricin Loop (SRL) of the ribosome, a process called depurination, which blocks the binding of elongation factors to the ribosome, leads to a halt of protein biosynthesis and thus causes cell death ^3,7^. A similar mechanism is used by Stx and related AB type RNA-*N-*glycosidase plant toxins, overall designated ribosome-inactivating toxins (RIPs), such as ricin from *Ricinus communis* and abrin from *Abrus precatorius*. However, those toxins engage other cellular receptors, especially with different oligosaccharide residues like *N*-acetylglucosamine- and galactose-containing receptors ^8^.

Although *N*-glycosydase activity is in general associated with RIPs, their effect on the SRL and shorter RNA or single-stranded DNA (ssDNA) SRL mimics, generated to facilitate detection of RIP activity, might be different. For example, ricin requires the recognition sequence GAGA as the minimal sequence for activity in rRNA ^9^ but saporin also depurinates adenine-containing 39-mer ssDNA substrates without the recognition sequence ^10^. In addition, Stx releases adenine from 10-mer rRNA SRL substrates including GAGA ^11^, however, it is not known whether short ssDNA SRL can serve as substrate.

More than ten years after the large STEC O104:H4 outbreak, timely and qualified detection of STEC including isolate recovery in patients, animals, and food remains of high importance but is still challenging, time-consuming, and labor-intense. Currently, detection of STEC is increasingly performed by means of molecular tools, such as Stx gene (*stx*) polymerase chain reaction (PCR), from stool or food matrices containing background flora, i.e. from strain mixtures, however often isolate recovery is not successful ^12^. Here, false-positive results may play a role because the PCR signal may be derived from non-vital bacteria or the free *stx* phage ^13^. Correlation of *stx* presence to a specific strain and accordingly isolate recovery is important for the following reasons: for strain risk profiling allowing disease outcome prediction, treatment considerations, decisions on quarantine regulations, or when detected in food for product withdrawal and for subsequent molecular epidemiological analysis permitting disease cluster and respective source recognition ^14^.

The so far published activity-based tests for *N-*glycosidase toxins include technically demanding, time-consuming, and cost-intense equipment or procedures, such as LC-MS/MS or multi-step enzymatic detection of the released adenine ^11,18–20^. Consequently, the availability of an easy-to-perform and cost-effective rapid test for identification of Stx and STEC on the isolate level would be of tremendous value for infection and pathogen diagnostics. In the here presented study, we developed a fluorescent enzyme substrate for rapid and simplified functional detection of Stx and STEC.

## Materials and Methods

### Experimental Design

In the here presented study, we developed a fluorescent enzyme substrate for rapid and simplified functional detection of Stx and STEC. We designed synthetic FRET-based ssDNA SRL substrates that conferred a fluorescence signal after cleavage by Stx. We further aimed to optimize the reaction conditions and in total validated the assay for 65 STEC strains and 11 *Shigella* strains, and 17 strains not producing Stx. Most of the strains originated from the collection of the German National Reference Centre (NRC) for Salmonella and other bacterial enteric pathogens of the Robert Koch-Institute.

### Bacterial strains, eukaryotic cell lines, growth conditions, and preparation of culture supernatants, and single colonies

Strains used in the study are listed in table S2 (STEC), table S3 (other *E. coli* pathovars without *stx*), table S4 (other enteropathogenic bacteria without *stx*), and table S5 (*Shigella*). A variety of strains were employed: 65 STEC strains harboring *stx1* and/or *stx2* incl. different *stx* subtypes (s*tx1a, stx1c, stx1d, stx2a-g*) and 30 different serotypes were analyzed (Tab. S2). Reference strains of STEC O157:H7 EDL933 harboring *stx1a* and *stx2a* and an isogenic *stx* negative knock out mutant EDL933 ∆*stx1/2* were used as controls (Tab. S2). Further strains, such as other intestinal pathogenic *E. coli* belonging to pathovars EPEC, EAEC, EIEC (Tab. S3), and other intestinal pathogenic bacteria, such as *Salmonella enterica, Yersinia enterocolitica*, all without *stx* (Tab. S4), and *Shigella* spp. with *stx1*a or *stx2a* were used (Tab. S5). All strains, excluding the reference strains, are clinical strains and were collected and characterized by the German National Reference Centre (NRC) for *Salmonella* and other Bacterial Enteric Pathogens.

All bacterial strains were streaked out and grown on LB agar overnight for single colonies at 37 °C or cultivated in LB broth (high NaCl 10g/L) for 18 h, 250 rpm, 37 °C. 12 ng/mL ciprofloxacin (Cip, Sigma-Aldrich, Darmstadt, Germany) was used in these media as an alternative for mitomycin C (MMC) for the induction of the Stx production ^21^. Culture supernatants were obtained by centrifugation (8,000 g, 10 min) and filtration (0.22 μm filters; Sartorius, Göttingen, Germany). Stx production was tested using Vero cell cytotoxicity assay (supplement), Enzyme Linked Immunosorbent Assay (ELISA) (supplement), and the here described Stx activity assay. Culture supernatants were stored at 4 °C for up to one week or at -20 °C for up to six months.

The Vero cell line (ACC33; German Collection of Microorganisms and Cell Cultures GmbH, Braunschweig, Germany) was cultured at 37 °C, 5 % CO_2_ in DMEM (Capricorn Scientific, Ebsdorfergrund, Germany) with 10 % Fetal Bovine Serum (FBS; Capricorn Scientific).

### *E. coli* serotyping and virulence gene analysis

The presence of *stx* was determined using colony PCR after growth on LB agar as described by Cebula et al. ^22^. Stx subtype and the presence of *eaeA* were determined by PCR or were extracted from the genome sequence ^4,23,24^. *E. coli* O and H antigens were determined by microtiter agglutination method or extraction from the genome sequence as described elsewhere ^24,25^.

### Enzyme Linked Immunosorbent Assay (ELISA)

The qualitative analysis of Stx in culture supernatants was performed using the commercially available Ridascreen Verotoxin Stx ELISA (r-Biopharm AG; Pfungstadt, Germany) as specified by the manufacturer. For the quantitative analysis, a custom-made Stx sandwich ELISA was used. Briefly, a 96 well plate (MaxiSorp; Nunc, Thermo Fisher Scientific, Germany) was coated with 10 μg/mL of a capture antibody in 50 μL Phosphate Buffered Saline (PBS) (13C4, hybridoma cell line from American Type Culture Collection, Manassas, USA, for Stx1; MBS311736, MyBioSource Inc., San Diego, USA, for Stx2) overnight at 4 °C and blocked with casein buffer (Senova, Jena, Germany) for 1 h at room temperature. After washing using PBS with 0.1 % Tween 20, 50 μL of diluted STEC culture supernatants or Stx standard dilution series were added and incubated for 2 h at room temperature. After washing, the bound Stx of STEC culture supernatants was detected using a biotinylated detection antibody (MBS311734 for Stx1, MyBioSource Inc., San Diego, USA; 11E10, hybridoma cell line from American Type Culture Collection, Manassas, USA; and BB12^27^, Toxin Technology Inc., Sarasota, USA, for Stx2) incubated for 1 h at room temperature. After washing, the ELISA was developed with PolyHRP40 (Senova GmbH; Jena, Germany) and substrate 3,3’,5,5’-tetramethybenzinine (TMB, SeramunBlau slow2 50, Seramun Diagnostika, Heidesee, Germany). The color reaction was stopped by 0.25 M sulfuric acid. After absorption detection at 450 nm versus 620 nm (Tecan Infinite M200), the concentration of Stx in culture supernatants was calculated using Stx1 and Stx2 standards (Toxin Technology Inc., USA) of known concentration in the range from 0.3 pg/mL to 100 ng/mL.

### StxSense SRL substrates for Stx detection

Four synthetic ssDNA substrates for Stx detection were designed based on the SRL sequence of *Rattus norvegicus* and commercially synthesized (Integrated DNA Technologies [idt], Leuven, Belgium; or biomers.net GmbH; Ulm, Germany). These substrates include the described SRL recognition sequence GAGA for RIPs and fluorophore / quencher pairs (Table 1). For all substrates, the quencher was at the 3’ end. **StxSense 1** contained a Cy5 fluorophore at the central adenine, which is specifically depurinated by Stx. All other substrates, substrates **StxSense 2 – StxSense 4**, were labeled with a 6-FAM fluorophore at the 5’ end. Substrates were synthesized by a commercial provider (e.g. integrated DNA technologies, idt) and quality control data for **StxSense 1 – StxSense 4** are shown in Fig. S2 to Fig. S5.

**Table 1.** Synthetic ssDNA substrates for the detection of Stx N-glycosidase activity. Studied SRL substrates and their respective sequences, features, fluorophores (F): Cy5 = Cy5 fluorophore; FAM = 6-FAM fluorophore (fluorescein) and quencher (Q). Sequence labeling: **bold** = recognition sequence including the adenine (<u>A</u>) targeted for depurination; blue and *italic* = modified ends sequence.

| substrate | sequence (5'→ 3') | k-mer | fluoro-<br>phore (F) | quencher<br>(Q) | characteristics |
| --- | --- | --- | --- | --- | --- |
| StxSense 1 | CTG AAC TCA GTA **CGA** (Cy5)<br>**GAG** GAA CC GTT CAG (Q) | 29 | Cy5 | BMN-Q620 | Adenine labeling |
| StxSense 2 | (FAM) TCA GTA **CGA** **GAG**<br>GAA CC (Q) | 17 | 6-FAM<br>(5' end) | BHQ-1<br>(3' end) | Reduced SRL sequence<br>compared to StxSense 1 |
| Stxsense 3 | (FAM) TCA GTA **CGA** **GAG**<br>**GAG** AGG AAC C (Q) | 21 |  |  | Double recognition<br>sequence **GAG**A |
| StxSense 4 | (FAM) *AC TTA* GTA **CGA** **GAG**<br>GA AC*AG T* (Q) | 21 |  |  | Modified ends before<br>F/Q |

### Stx *N-*glycosidase enzyme assay using culture supernatants and single colonies

StxSense synthetic ssDNA substrates that are fluorophore-coupled (Tab. 1) were used to detect the *N-*glycosidase activity of Stx. For culture supernatant samples, 14.6 μL of 100 mM ammonium acetate (Fluka, item no. 09690, VWR International LLC, Pennsylvania, USA), adjusted to pH 4 with HCl (32 %; item no. 4625.1, Carl Roth GmbH, Karlsruhe, Germany) and 0.4 μL of the substrate (100 mM stock in dH_2_O; 2 μM final test concentration) were mixed (reaction mix) and transferred into a white 96 well plate (Eppendorf Twin-Tec, item no. 0030132718, Eppendorf SE; Hamburg, Germany). For single colony samples, 19.5 μL 10 mM ammonium acetate, adjusted to pH 4 with HCl, and 0.5 μL of the substrate (100 mM in dH_2_O; 2 μM final concentration) were mixed (reaction mix) and transferred into a white 96 well plate. Subsequently, 5 μL of the culture supernatant or a single colony prepared as described above was added. The assay was carried out using the following parameters: 44 °C, 1 h to 12 h reaction time, detection of fluorescence, for **StxSense 1** Cy5 or **StxSense 2 – StxSense 4** FAM filter, at various time points using a real-time instrument (CFX96; Bio-Rad Laboratories, California, USA). Stx activity was analyzed by depicting the Relative Fluorescence Units (RFU) versus time graphically in GraphPad Prism (GraphPad Software; San Diego, USA).

### Specificity of Stx detection

The Stx enzyme assay was examined by using STEC strains expressing *stx1* and/or *stx2* as well as the *stx* subtypes (*stx1a, c, d, stx2a-g*). In addition to strains of the serotype O157:H7, strains of 29 additional STEC serotypes were analyzed. To exclude cross reactivity, other intestinal pathogenic *E. coli* (EAEC, EPEC, EIEC) and other intestinal pathogenic bacteria (such as *Yersinia, Salmonella*) without *stx* were tested.

### Sensitivity of Stx detection

The limit of detection (LOD) was calculated using quantified culture supernatants (see section ELISA). For STEC culture supernatants comprising Stx subtypes Stx1a and Stx2a, a 1:2 dilution series in the range of 1 ng/mL to 126 ng/mL (Stx1a) or 138 ng/mL (Stx2a), respectively, was prepared and 5 μL of each dilution step were analyzed for enzyme activity. The LOD was determined from regression curves using GraphPad Prism and the equation LOD (RFU) = 3 · SD/b, in which LOD is limit of detection, SD is standard deviation and b is the slope of the regression curve.

### Data analysis

Values of the enzyme assay curves were obtained with CFX Maestro (Bio-Rad Laboratories, Inc.), and were illustrated and analyzed using GraphPad Prism (GraphPad Software; San Diego, USA). Statistical analysis was performed for the RFU values at 2 h, 4 h, 8 h, or/and 12 h. All experiments were examined for normal distribution. Normally distributed values were statistically analyzed using double-sided t test, corrected with Welch, whereas for non-normally distributed values the Mann-Whitney-U-test was used (see supplement).

### Graphical representation

Graphs were created using GraphPad Prism, version 9.10.0.22 (GraphPad Software, LLC), and figures using BioRender.com (Toronto, Canada).

## Results

### Principle of the enzymatic STEC detection assay based on Stx *N-*glycosidase activity

We aimed to detect Stx *N-*glycosidase activity by employing synthetic SRL mimics equipped with a fluorophore (e.g. 6-FAM) and quencher (Q) pair which, when in physical proximity, result in FRET-based quenching, i.e. no fluorescence (Fig. 1A) ^28^. In the presence of Stx, the SRL is depurinated, resulting in a cleavage of the sugar phosphate backbone. When the fluorophore and the quencher are attached to positions down- and upstream of the depurinated site, the loss in physical proximity upon cleavage induces a rapid increase of a fluorescent signal indicative of Stx (Fig. 1A). The main steps of the envisioned STEC detection assay include 1) strain cultivation, 2) generation of bacterial supernatants or single colonies, 3) preparation of the reaction mix, 4) reaction plate preparation and reaction start, and 5) fluorescence detection as depicted in Fig. 1B.

**Figure 1.**
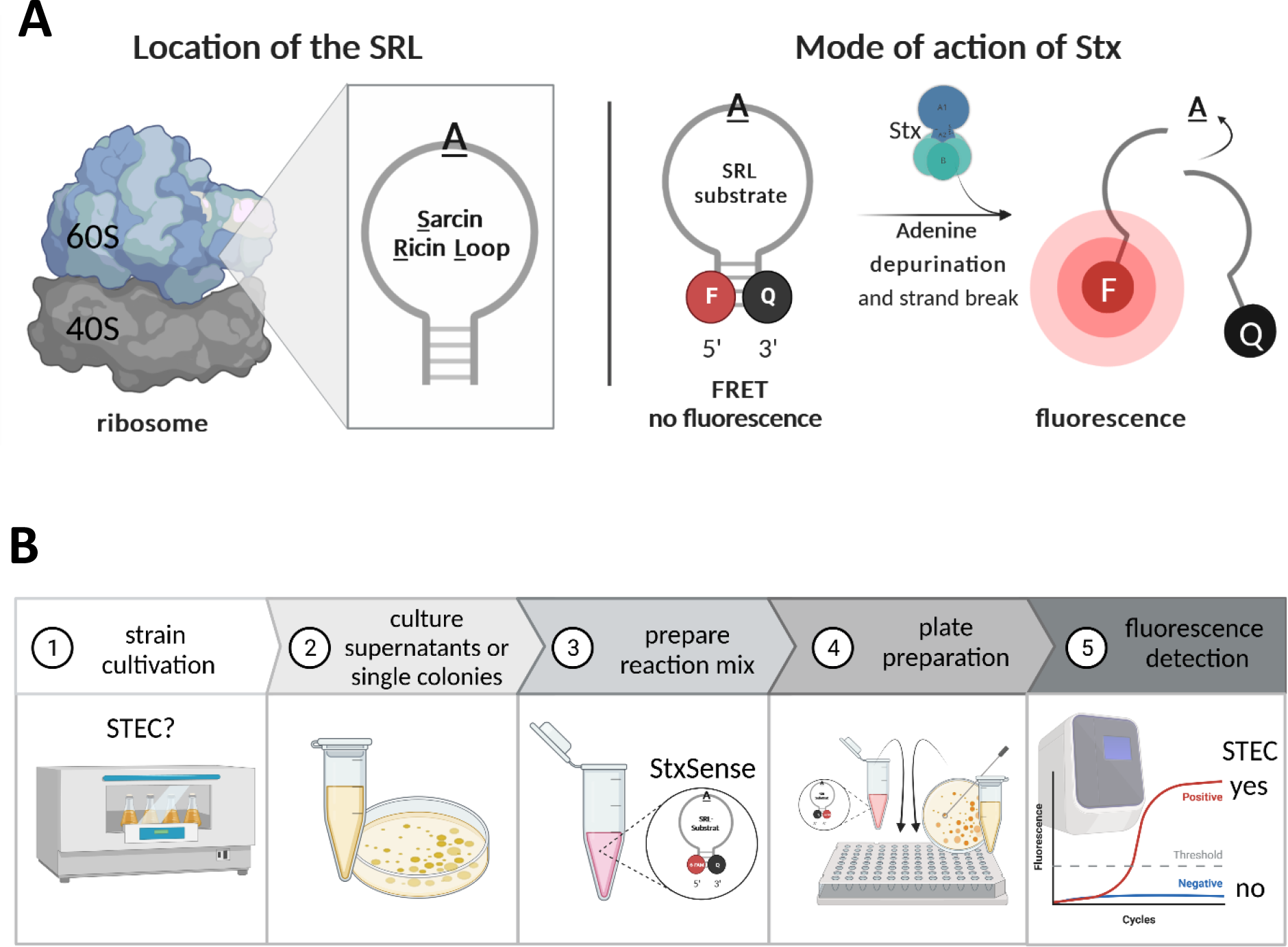
Principle of the enzymatic assay for STEC detection based on Stx *N*-glycosidase activity. **(A)** Location of the Sarcin Ricin Loop (SRL) targeted by Stx activity within the 60S subunit of the ribosomes. A synthetic SRL mimic with fluorophore (F) and quencher (Q) is used *in vitro* to detect the enzymatic activity of Stx. In presence of Stx, the SRL adenine is depurinated and the sugar phosphate backbone is cleaved. As a result, the fluorophore and quencher are no longer in physical closeness resulting in a fluorescent signal. **(B)** Main steps of STEC detection assay.

### Design of fluorescently labeled ssDNA Stx substrates based on the SRL

In order to increase the stability of the reagent, an ssDNA template was chosen instead of the natural Stx RNA-based substrate. We designed four ssDNA substrates **StxSense 1** to **StxSense 4** of different lengths (17 to 29 nt) based on the SRL sequence from *Rattus norvegicus* (Fig. 2A, Tab. 1). **StxSense 1, 2**, and **4** included one recognition sequence G<u>A</u>GA (target adenine for depurination underlined), while **StxSense 3** contained two of these to promote the chance of cleavage ^29^. Cy5 and 6-FAM were used as fluorescence markers; the first was coupled to the targeted adenine in **StxSense 1** and the latter was located at the 5’ end of the substrates in **StxSense 2** to **StxSense 4**. Quencher BMN-Q620 was used in **StxSense 1** and was located within the substrate sequence, whereas quencher BHQ-1 was located at the 3’ end in substrates **StxSense 2** to **4**. Further, **StxSense 4** comprised adapted sequence ends (5’ ACTT and 3’ AGT) to potentially improve the fluorescence signal after cleavage. Further, the base pairing of the two most distal bases brings fluorophore and quencher closer together, so that the quenching effect would be optimal (Fig. 2A).

**Figure 2.**
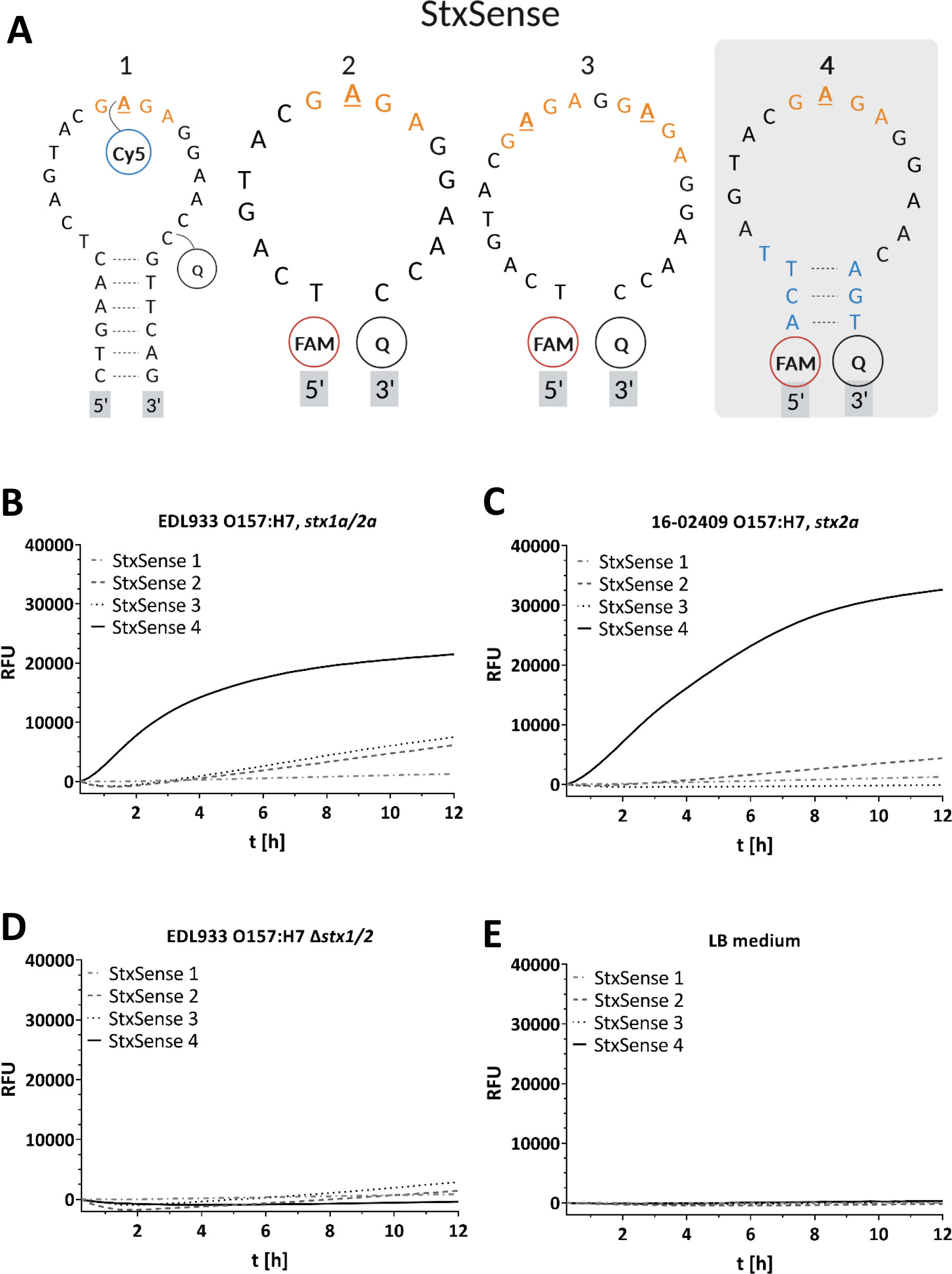
Fluorescently labeled ssDNA substrates based on the SRL detect Stx. **(A)** Different synthetic SRL substrates StxSense 1-4. Cy5 and FAM (here 6-FAM) denote fluorescence markers and Q a quencher (BMN-Q620 in StxSense 1 and BHQ-1 in StxSense 2-4). Stx recognition sequence is highlighted in orange embedded within the SRL sequence of *Rattus norvegicus* and the target adenine for depurination is underlined. StxSense 3 contains two recognition sequences. StxSense 4 contains adapted sequence ends (5’ ACTT and 3’ AGT) compared to the wildtype sequence of *Rattus norvegicus* to improve the fluorescence signal. **(B)-(E)** Detected fluorescence as a marker of substrate hydrolysis by Stx from positive control EDL933 O157:H7, test strain STEC 16-02409 O157:H7, and negative controls EDL933 O157:H7 ∆*stx1/2* and LB medium. For both Stx producing strains (EDL933 O157:H7 and 16-02409 O157:H7), substrate StxSense 4 was the optimal substrate for detection of Stx activity. The results represent the medians of triplicate samples (n = 3) and are representative of three independent experiments. Statistical analysis was performed by unpaired, double-sided t test (*, p < 0.05; **, p < 0.01; ***, p < 0.001), with results compared to those of the EDL933 O157:H7 ∆*stx1/2*. For statistics and Vero cell cytotoxicity assay see Fig. S6. RFU, relative fluorescence units; t [h], time [hours].

### Substrate StxSense 4 showed highest Stx-dependent fluorescence using STEC culture supernatants

To test the designed substrates for Stx detection, culture supernatants of two STEC strains, specifically, reference strain STEC O157:H7 EDL933 comprising *stx1a* and *stx2a* and clinical STEC O157:H7 isolate 16-02409 comprising *stx2a* were incubated with 2 μM substrates **StxSense 1** to **StxSense 4** in depurination buffer at pH 4 and at 44 °C for up to 12 h. The fluorescence readouts were compared to *stx1/2*-deficient EDL933 ∆*stx1/2* strain and LB medium, which served as negative controls. For both STEC strains, substrate **StxSense 4** induced the highest fluorescence readout among the four substrates from ∼30 min to 1 h (Fig. 2B and C, statistics Fig. S6A and S6B), whereas the negative controls did not show substantial fluorescence over the whole time period for **StxSense 4** (Fig. 2D and E, statistics Fig. S6C and S6D). Further, we tested other intestinal *E. coli* pathovars and other intestinal pathogens without *stx* for assay interference. EPEC, EAEC, and EIEC strains (Fig. S6E, statistics Fig. S6F) and additional strains of different *Salmonella enterica* serovars and of *Yersinia enterocolitica* (Fig. S6G, statistics Fig. S6H) did not show cross-reactivity. Additionally, the tested strains behaved as expected in Vero cell cytotoxicity assays which is an established method for Stx detection. Specifically, only strains comprising *stx*, such as STEC strain EDL933, showed reduced cell viability, but not EDL ∆*stx1/2* or other intestinal pathogenic bacteria all without *stx* (Fig. S6I and J). In conclusion, the experiments indicated that the substrate **StxSense 4** was most suited for STEC and Stx detection and that other Stx-negative intestinal pathogenic bacteria did not show cross-reactivity. Therefore, **StxSense 4** was used for all further experiments in the study.

### Optimal assay conditions for Stx detection in culture supernatants

We optimized important assay parameters, such as substrate concentration, concentration of the depurination buffer ammonium acetate, the nature of the 96 well plates, and reaction temperature. First, the **StxSense 4** concentration was analyzed in the range of 1 μM to 8 μM. For robust Stx detection from STEC EDL933 culture supernatants within 30 min/1 h, 2 μM **StxSense 4** were sufficient (Fig. S7A and statistics S7B). Importantly, fluorescence readout at earlier time points further increased especially to a concentration of 5 μM. Second, the depurination buffer ammonium acetate (pH 4) was applied in 10 mM and 100 mM concentrations. A 100 mM ammonium acetate buffer led to higher readouts for STEC strains EDL933 and 16-02409 (Fig. S7C and statistics S7D). Third, the nature of the 96 well plates had an influence on Stx detection. For both STEC culture supernatants, the white plates performed better than transparent ones (Fig. S7E and statistics S7F). Fourth, the influence of the reaction temperature was analyzed in the range of 37 °C to 45 °C. For STEC EDL933 culture supernatants, the fluorescence readout was highest from ∼44 °C, but substantial readouts were already observed from 37 °C (Fig. S7G and statistics S7H). To account for both maximal signal as well as cost-effectiveness, the following assay parameters were defined for STEC culture supernatants and used throughout the study: 2 μM **StxSense 4**, 100 mM ammonium acetate, white 96 well plates, and reaction temperature of 44 °C.

### The limit of detection (LOD) for the assay lies in the range of ELISA

We further analyzed the limit of detection of this assay. To that end, we used culture supernatants of a solely Stx1a- and another solely Stx2a-producing strain, O111:H8 20-01044 and O157:H7 16-02409, respectively. The culture supernatants were diluted and the Stx concentrations were determined by sandwich ELISA; in parallel RFU readout was analyzed by means of the novel assay. LOD for Stx1a was determined as 10 ng/mL (Fig. 3A) and LOD for Stx2a was 29 ng/mL (Fig. 3B). Those LODs are comparable to other standard Stx tests, such as ELISA and Vero cell cytotoxicity tests where LOD ranges from 10 pg/mL to 32 ng/mL ^17^.

**Figure 3.**
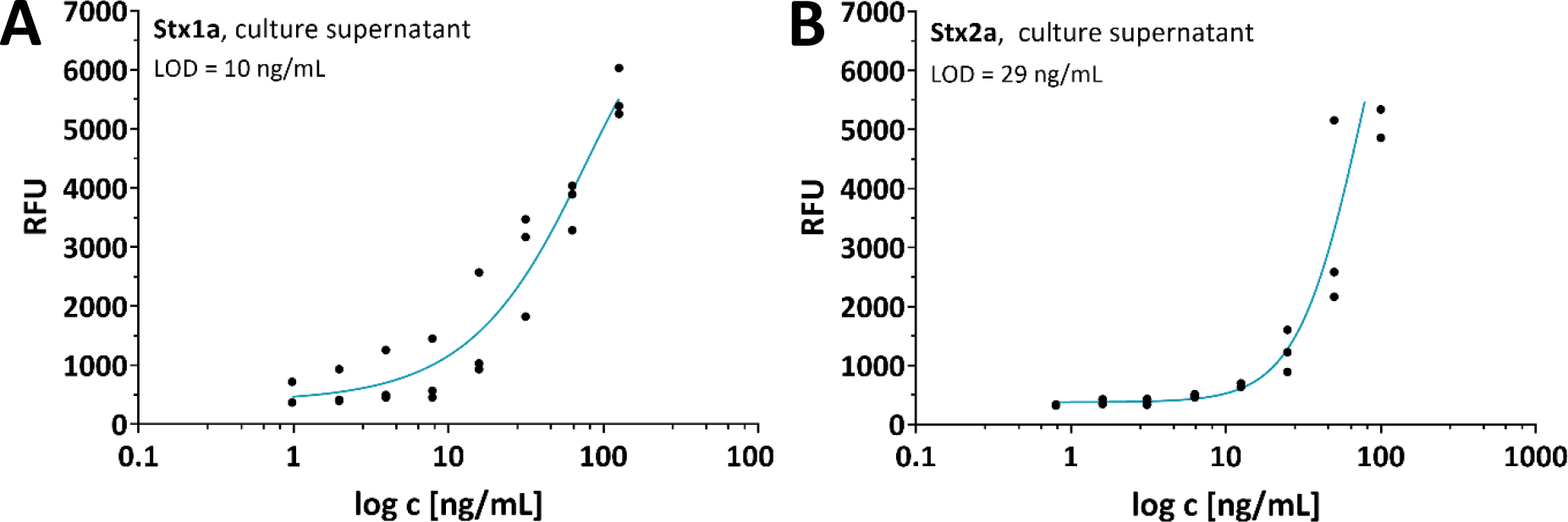
Limit of detection correlation analysis for STEC Stx1a and Stx2a. Concentration-dependent analysis of Stx activity in 100 mM ammonium acetate, pH 4, 44 °C of strains 20-01044 O111:H8, *stx1a* and 16-02409 O157:H7, *stx2a*. Plotted are the log c [ng/mL] of the Stx ELISA quantified culture supernatants versus the detected RFU. The correlation was obtained using an exponential function and equation LOD = 3 * SD / b and LOD is limit of detection, SD is standard deviation and b is slope of regression curve. Shown is the representative plot of one determination in triplicates from a total of three representative independent determinations (n = 3). RFU, Relative Fluorescence Units.

### Detection of Stx1 and Stx2 subtypes across different STEC serotypes and *Shigella* spp

Three different subtypes of *stx1* and fourteen subtypes of *stx2* have been described ^4,5^. Therefore, we analyzed whether the most relevant Stx subtypes are detected by the assay. To this end, additional STEC strains of serotypes harboring *stx1a, stx1c, stx1d* or/and *stx2a* to *stx2g* were analyzed using culture supernatants or single colonies. The latter was additionally implemented because the release of Stx1 into culture supernatants is lower compared to Stx2 ^30^. In culture supernatants, strains showing Stx2a, Stx2b, a combination of Stx1a and Stx2a, Stx2f, Stx2g, Stx2d, and Stx2c were detected between 30 min and 2 h reaction time, whereas the ones with other Stx types, including Stx2e, and Stx1a, required more time (Fig. 4A, statistics Fig. S8A). A weak signal was observed for Stx1c and Stx1d after >10 h (Fig. 4A, statistics Fig. S8A and S8B). All tested *stx* positive strains revealed Vero cell toxicity as a standard measure of Stx (Fig. S8C). Using single colonies of the same strains instead of culture supernatants, especially the combination of Stx1a and Stx2a, several Stx2 (Stx2g, Stx2f, Stx2d), and after ∼3-4h Stx1a, Stx1c and Stx1d and some other Stx2 were detected, but also the background of strain EDL933 ∆*stx1/2* rose (Fig. 4B, statistics Fig. S4D). When the assay was performed with single colonies, we noted that 10 mM ammonium acetate buffer instead of 100 mM buffer produced higher fluorescence signals and therefore 10 mM buffer was used for single colony analysis (Fig. S4E and statistics S4F).

**Figure 4.**
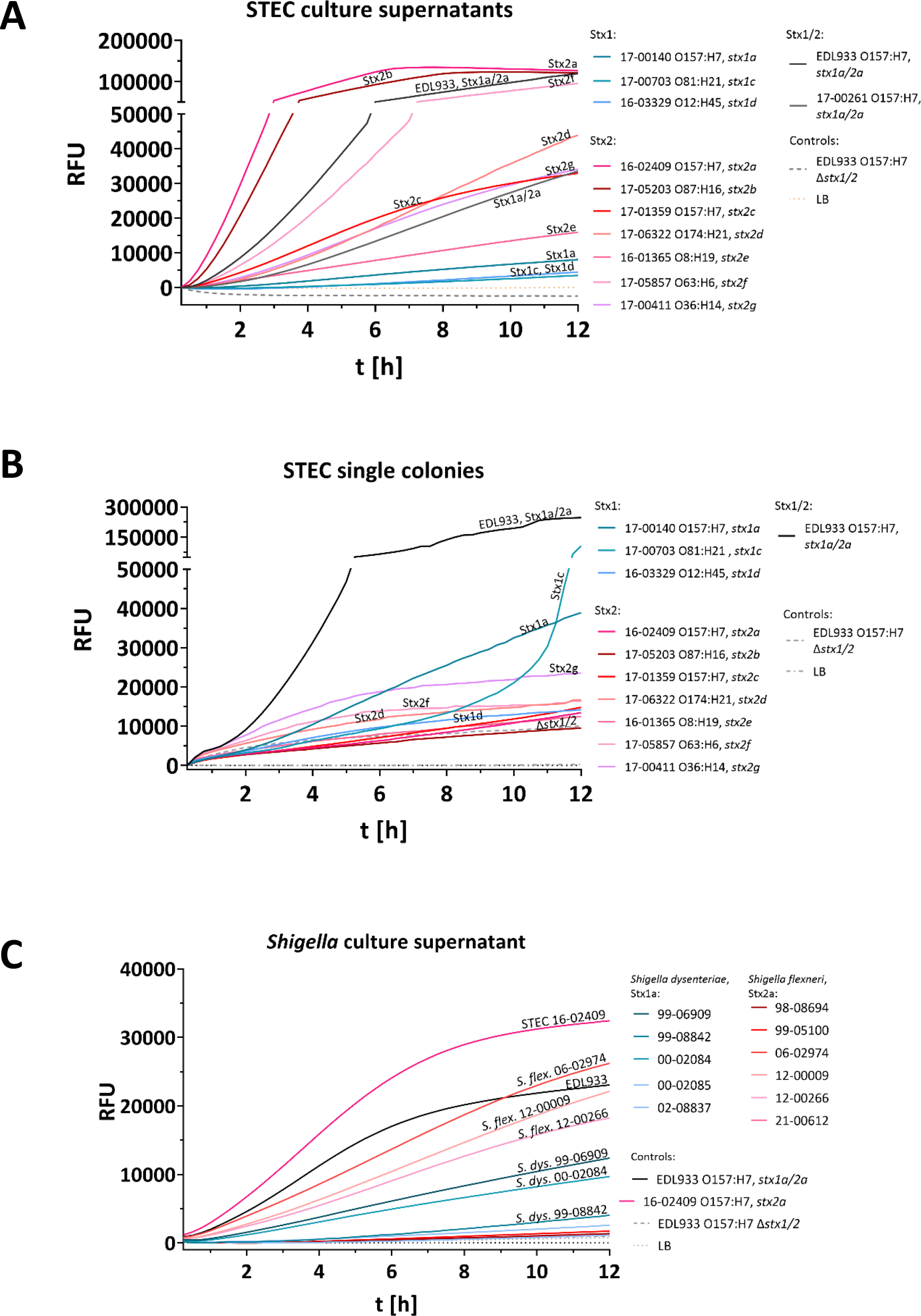
Detection of relevant Stx1 and Stx2 subtypes was possible in a variety of STEC serotypes and *Shigella* spp. Detected fluorescence as a marker of substrate hydrolysis by Stx from STEC culture supernatants and single colonies. **(A)** The assay detects Stx2 subtypes and some Stx1 subtypes when culture supernatants of bacteria grown in LB medium supplemented with 12 ng/mL ciprofloxacin are used. Since Stx1 is scarcely released into the culture supernatant, detection of Stx1 in culture supernatants is strain dependent. **(B)** Using single colonies grown on LB agar supplemented with 12 ng/mL ciprofloxacin, Stx1 and Stx2-producing STEC strains are detected within 10 h. **(C)** *Shigella flexneri* strains releasing Stx into culture supernatants are detected by the assay. The results represent the medians of triplicate samples (n=3) and are representative of three independent experiments. Statistical analysis was performed by Mann-Whitney test (A, B) for non-normally distributed samples and unpaired, double-sided t test (C) for normally distributed samples (*, p < 0.05; **, p < 0.01; ***, p < 0.001), with results compared to those of the EDL933 O157:H7 ∆*stx1/2* (negative control; -- [grey dashed lines]). For statistics and Vero cell cytotoxicity assay see Fig. S8 and S9. RFU, relative fluorescence units; t [h], time [hours].

Various clinical STEC strains of frequently disease-associated O types in addition to O157, such as O26, O91, O111, O113, O121, O145 all resulted in a positive Stx assay signal from culture supernatants (Fig. S8G, statistics S8H, and Vero cell assay S8I). Further, 46 additional clinical STEC strains from the NRC collection of a variety of serotypes and *stx* subtypes were tested based on culture supernatants and were all detected by the Stx enzymatic assay, except six strains did not result in a significant RFU change (five strains showing *stx2e* and one *stx2d*) and correspondingly did not reveal cytotoxicity (Fig. S8J, L, N; statistics Fig. S8 K, M, O, cytotoxicity Fig. S8P).

Further, some *Shigella* strains may harbor *stx* ^1^. Indeed, *S. flexneri* strains releasing Stx2*a* into the culture supernatant, such as strains 06-02974, 12-00009, and 12-00266, or *S. dysenteriae* showing Stx1a, such as strains 99-06909, and 99-08842, 00-02084, resulted in a fluorescence signal, conferred cytotoxicity towards Vero cells, and resulted in Stx detection by Stx ELISA (statistics Fig. S9A and B, cytotoxicity Fig. S9C, and Stx ELISA Fig. S9D). As expected, the five *Shigella* strains which did not result in a fluorescent signal, did not show cytotoxicity towards Vero cells and three of the six did not release Stx into the culture supernatant (Fig. S9A to D).

In conclusion, standard assay conditions allowed for detection of all 59 STEC strains producing Vero cell cytotoxicity, comprising the manifold Stx1 and Stx2 subtypes, and a wide variety of different STEC serotypes and also *Shigella* spp. Further, implementation of culture supernatants is optimal for Stx2-producing strains, and single colony analysis can improve the detection of Stx1-producing strains.

## Discussion

Timely and qualified detection of STEC including isolate recovery is of high importance but still challenging and labor-intense. Current PCR procedures target *stx*, but the isolation of STEC from the PCR-positive samples succeeds in only about 42 % of cases ^12^. Although PCR methods for determination of *stx* subtypes correlating to disease severity are available ^31^, they do not allow conclusions on whether active Stx is produced and on Stx amounts; two important parameters which might be linked to the fate of disease and progression into HUS. In addition, other tests targeting resistance or metabolic features are only covering a subset of the multi-faceted STEC group. Thus, an easy-to-perform rapid test for all STEC represents an unmet diagnostic need. Therefore, we developed an STEC detection method based on the *N-*glycosidase activity of Stx.

To that means, we designed different synthetic FRET substrates mimicking the SRL which after cleavage by Stx result in a real-time fluorescence signal (Fig. 1A and 2A). Substrate **StxSense 4** yielded in the highest fluorescent readout (Fig. 2B and 2C). The other substrates did either not respond to Stx at all, such as **StxSense 1**, containing the fluorescence marker at the target adenine for depurination, or revealed a significantly lower fluorescence signal than **StxSense 4**. These, **StxSense 2** and **StxSense 3**, like the optimal substrate **StxSense 4**, had the fluorescence marker at the 5’ end and the quencher on the 3’ end. Differences compared to **StxSense 4** were that in **StxSense 2** and **StxSense3** no modified ends were added. In **StxSense 3**, an additional G<u>A</u>GA recognition sequence was added which however did not lead to improved activity. Therefore, we conclude substrate **StxSense 4** was optimal which contained a central recognition site G<u>A</u>GA, the fluorescence marker and quencher at the distal ends of the substrate flanked by modified ends.

Previous work on RIP substrates, including Stx substrates, were not linked to a fluorescence marker and encompassed typically RNA and for plant RIPs in some cases ssDNA ^20^. The sequences mostly included the G<u>A</u>GA recognition site and varied in length between 10 and 29 bases. Often, release of adenine was quantified by means of enzymatic ADP to ATP conversion or liquid chromatography coupled mass spectrometry (LC-MS).

In a 2013 patent (No.: US 10,907,193 B2), a fluorescence-labeled ssDNA substrate was used to detect the activity of ricin in a real-time approach. Ricin depurinates the specific adenine of the recognition sequence in a synthetic ssDNA substrate and creates an abasic site. By adding a lyase, the DNA is cleaved at this site and accordingly a fluorescence signal is generated. It seems that the strand break might be a limiting factor of some plant RIPs requiring the addition of strand-breaking enzymes. This is not the case for Stx detection, as shown in our study. The substrates used in the patent varied in length from 26 to 31 nt and contained the fluorescent group at the 5’ and the quencher at the 3’ end. It was concluded that both shape and sequence of the substrates, some of which did not harbor the G<u>A</u>GA recognition sequence because certain plant toxins might depurinate any adenine in the sequence ^10^, seem to play a role in catalysis by ricin.

The here introduced assay requires minimal efforts (Fig. 1B and Fig. 5) and is as cost-effective as PCR methods. Specifically, bacterial culture supernatants or single colonies are directly incubated with reaction buffer composed of the pH 4 depurination buffer and the **StxSense 4** enzyme substrate. The novel assay importantly only requires the fluorescent SRL substrate which can be easily obtained from a commercial oligo/probe synthesizing company. However, for PCR approaches primers, dNTPs, and a polymerase and for qPCR additional DNA-binding dyes or a fluorescent probe are necessary.

**Figure 5.**
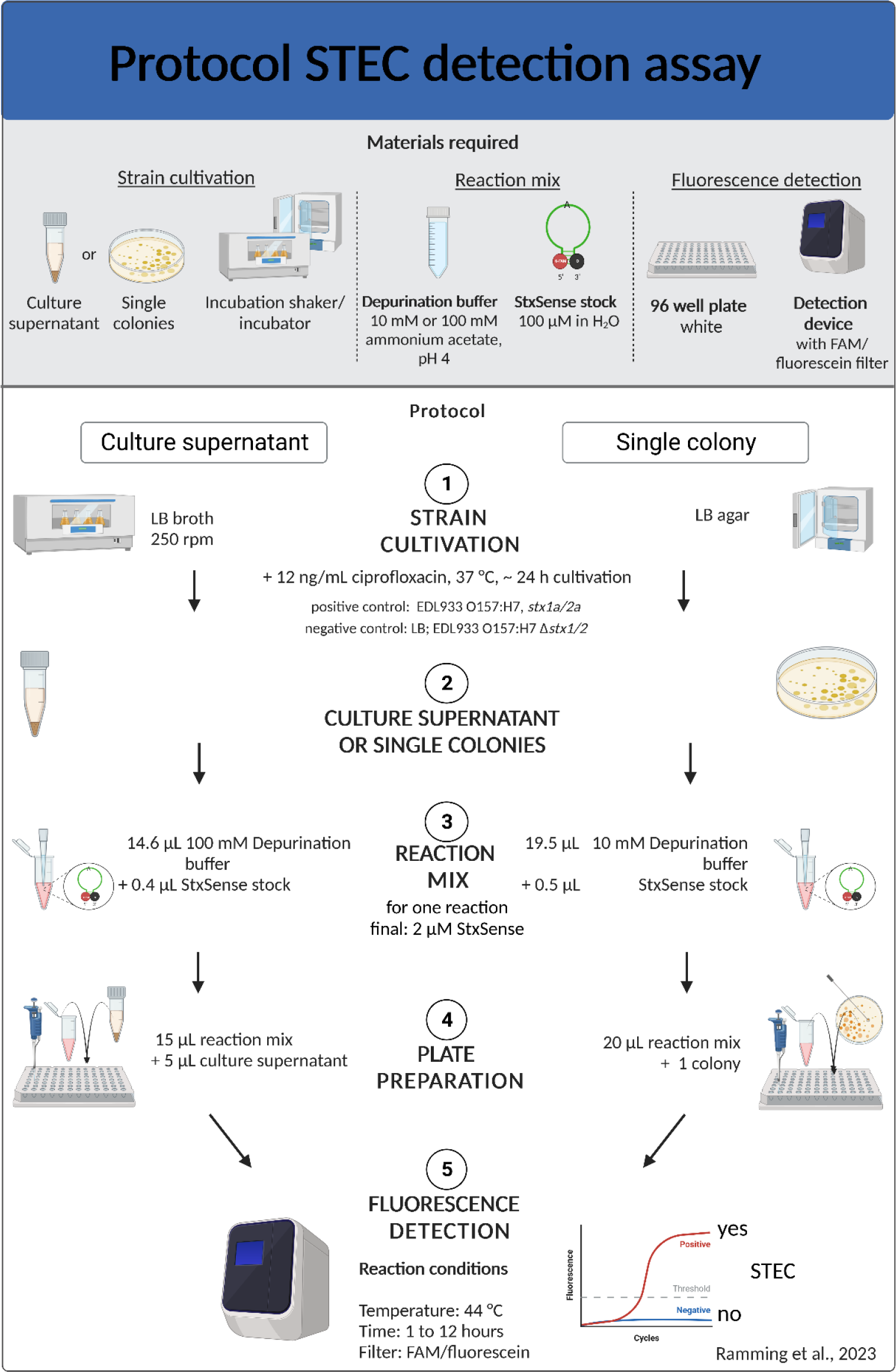
Assay protocol for the STEC detection assay. Overview of the materials required and the steps involved in the enzymatic assay for Shiga toxin detection from culture supernatants or single colonies. Briefly, after standard cultivation of STEC in LB broth or on LB agar, both with 12 ng/mL ciprofloxacin, the culture supernatant or single colonies are added to the reaction mix (ammonium acetate buffer pH4 containing SRL substrate) in a well of a 96 well plate. The enzymatic reaction is performed at 44 °C for up to 12 hours and the fluorescent signal (RFU, Relative Fluorescent Unit) is read with a detection device containing the appropriate FAM/fluorescein filter, such as a real time cycler or fluorescence gel imager. The supernatant or single colonies of EDL933 O157:H7, *stx1a/2a* are used as a positive control; LB, *E. coli* C600, or EDL933 O157:H7 ∆*stx1/*2 as negative controls defining the threshold.

Li and Tumer ^11^ used 2 μM substrate concentration for the detection of Stx activity. Here, Stx activity was determined by depurination of a 10-mer RNA substrate, further enzymatic conversion of the released adenine into ATP, and detection of a luminescent readout. A minimal substrate concentration of 2 μM was also found in our experiments for Stx detection, and fluorescence intensity increased for the tested reference strains up to a substrate concentration of 5 μM (Fig. S7A). For example, when 2 μM **StxSense 4** substrate were used, costs are ∼0.20 Euro/sample or ∼0.50 Euro/sample for 5 μM substrate concentration, respectively. Other materials required for the depurination buffer and 96 well plates sum up to less than 0.10 Euro/sample. Therefore, costs for the here described assay are in the range or even below PCR and qPCR ^32^. Furthermore, the LOD lies in the range of ∼10 to ∼30 ng/mL (Fig. 3A and B) and is therefore comparable to established methods, such as ELISA LOD of ∼30 ng/mL ^33^ or Vero cell cytotoxicity assay LOD of ∼10-32 ng/mL ^34^. Therefore, the combination of simplicity, cost-effectiveness, and sensitivity are clear advantages of the novel FRET-based assay.

Our data show that higher temperatures, e.g. 44 °C, were optimal for the assay readout. This observation has been described before for Stx and other RIPs, and even strand breaks after substrate depurination by plant toxins are further promoted by final short-term incubations at 90 °C ^29^. Additionally, acidic pH and a combination with higher temperatures is also beneficial for Stx enzymatic assays ^35^. This may be due to structural reorganizations occurring in Stx which open the catalytic site for better substrate access ^11^. Nevertheless, it is possible to perform the assay at 37 °C without a major reduction in fluorescence intensity (Fig. S7G and H). This fact might be important for a possible future adaptation of the assay as an agar plate-based STEC growth and detection medium.

In our study, we analyzed a total of 94 different strains including 65 STEC and 11 *Shigella* strains. All strains producing functional Stx in the detection sample yielded in Stx activity in the novel assay. Therefore, the assay can promote analysis of STEC and in addition may allow faster detection of relevant STEC strains, particularly such strains producing cytotoxicity and such predominantly associated with HUS development. Indeed, our data highlight high and consistent fluorescence readouts for Stx subtypes Stx2a, Stx2c, and Stx2d known for their superior HUS association ^36^. Interestingly, also STEC producing Stx2b showed high fluorescence readouts which may correlate to recent EFSA data about prolonged course of disease for such infections ^37^.

In addition to facilitating STEC detection, our assay provides an alternative method for analysis of Stx enzyme activity. So far, methods including 1) Stx toxicity towards susceptible eukaryotic cells, especially Vero cells, 2) ribosome inactivation in a cell-free *E. coli* protein synthesis system, or 3) depurination of SRL mimics without fluorescence marker, as outlined above, are used. The methods mostly require specialized equipment, such as mass spectrometers, eukaryotic cell culture, expensive reagents, or ribosome or sensitive DNA/RNA preparations ^18^. All these approaches are more complex and laborious than the method introduced here. The new method therefore might open up alternative test strategies for novel therapeutic approaches. Currently, there is no effective treatment for HUS, and only supportive care, such as fluid volume management, is recommended ^38^. Applying the novel Stx enzyme assay, antimicrobials can now be easily tested for their potential of Stx phage induction and accordingly therapy-associated increased Stx production ^21^. Further, the novel method may facilitate screens for substances inhibiting Stx activity.

## Supporting information

Supplemental Data_Ramming et al_2023

## Data availability

The data underlying this article are available in the article and in its online supplementary data.

## Acknowledgments

We thank Ute Strutz, Christian Galisch, Claudia Lampel, Karsten Großhennig, Petra Hahs, Susanne Karste from the Robert Koch-Institute for excellent technical assistance.

## Author contributions

I.R., B.G.D., M.B. and A.F. designed research; I.R., C.L., S.H., M.K., S.W., C.P., and A.Fr. performed research; I.R., C.L., S.H., M.K., S.W., C.P., A.Fr., B.G.D., M.B. and A.F. analyzed data; and I.R. and A.F. wrote the paper.

