## Supplemental Data_Ramming et al_2023 for "Rapid enzymatic detection of Shigatoxin-producing *E. coli* using fluorescence-labeled oligonucleotide substrates"

### Ramming et al., Tab. S1

**Tab. S1. Characteristics of the *Shigella* and STEC $AB_5$ Shiga toxins**

| | Stx | Stx1 | Stx2 | Reference |
|---|---|---|---|---|
| **Organism** | *Shigella dysenteriae* | STEC | STEC | |
| **Subtypes** | 1a | 1a, 1c, 1d | 2a-o | Scheutz et al., 2012<br>Yang et al., 2022 |
| **Protein sequence identity to Stx of *S. dysenteriae*** | 100 % | 91 – 99 %* | ~ 55 %* | Bergan et al., 2012 |
| **No. of amino acids** | | A1: 251<br>A2: 42 | A1: 250<br>A2: 47 | Li & Tumer, 2017 |
| **Active site in StxA1** | Glu 167 | Glu 167 covered by the A2 chain in the holotoxin (A1-A2) | Glu 166<br>open conformation in the holotoxin (A1-A2) | Yamasaki et al., 1991<br>Steyert et al., 2012<br>Jackson, 1990 |
| **Bacterial growth media and Stx production** | n.k. | Culture media affect bacterial growth and concentration of Stx in supernatants | | Rocha & Piazza, 2007 |
| **Activity of StxA1 on SRL RNA** | n.k. | $k_{cat}$ = 21.5 $min^{-1}$; lower compared to Stx2A1 | $k_{cat}$ = 62.6 $min^{-1}$; higher compared to Stx1A1 | Basu et al., 2015 |
| **Optimal pH for activity** | n.k. | *in vitro*: acidic pH (pH 4.5) | | Basu et al., 2015 |
| **Activity and temperature** | Activity loss<br>≥ 65°C | Activity loss<br>≥ 65°C | Activity loss<br>≥ 85°C | Chan & Ng, 2016 |

* Sequence identities of all Stx subtypes are summarized in Bergan *et al.,* 2012; n.k., not known

### Ramming et al., Fig. S1

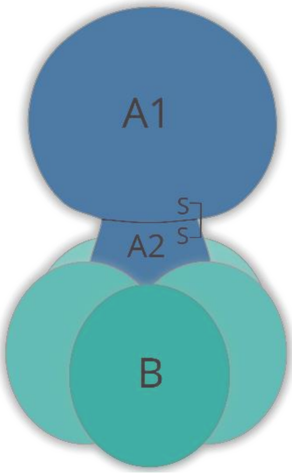


| Sub-unit | MW [kDa] | No. of amino acids (without signal peptide) | | Function | Reference |
|---|---|---|---|---|---|
| | | Stx1 | Stx2 | | |
| **A** | 32 | 293 | 297 | A1 and A2 connected by disulfide bridge after cleavage by furin | Gyles et al., 1988<br>Scheutz et al., 2012 |
| **A1** | 27.5 | 251 | 250 | Enzymatic activity, 28S rRNA *N*-glycosidase activity | Scheutz et al., 2012 |
| **A2** | 4.5 | 42 | 47 | Indispensable for holotoxin structure | Menge, 2020 |
| **B** | 7.7 | 69 | 68–70 | Binding to Gb3/CD77 receptor | Takeda et al., 1993 |

**Fig. S1. Structure of the STEC AB5 toxin Shiga toxin (Bergan et al., 2012) and characteristics of Stx1 and Stx2 A and B subunits.**

### Ramming et al., Fig. S2

**SRL-dA-Cy5**
00277426_1 DNA

02129832

5'-tca gta cg5 gag gaa cc-3'

| | | | | | |
|---|---|---|---|---|---|
| **Scale** | S | **Tm** | - | **Menge** | 3,0 OD |
| **Reinigung** | HPLC | **GC-Gehalt** | 52,00% | | 13,0 nmol |
| | Getrocknet | **Ext.Koeff.:** | 232300 | | 81,3 µg |
| **Länge** | 17 | A:5,0 C:4,0 | | **Vol.f.100pmol/µl** | 130 µl |
| **MW Soll** | 6275 g/mol | G:5,0 T/U:2,0 | | **Konz.** | - pmol/µl |
| **MW Ist** | 6277 g/mol | | | **Gelöst in** | - µl |

| | | | |
|---|---|---|---|
| **5'-Mod.:** | Keine | **Interne Mod. 5:** | dA-Cyanine 5 |
| **3'-Mod:** | BMN-Q620 | **6:** | Keine |
| | | **7:** | Keine |
| | | **8:** | Keine |

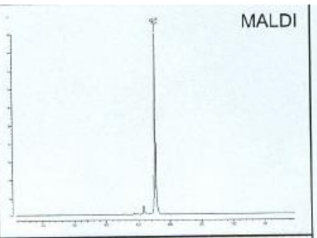


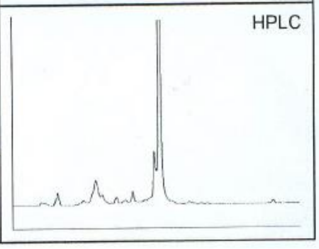


**Fig. S2. Quality control analysis report of StxSense 1 (biomers.net).**

### Ramming et al., Fig. S3

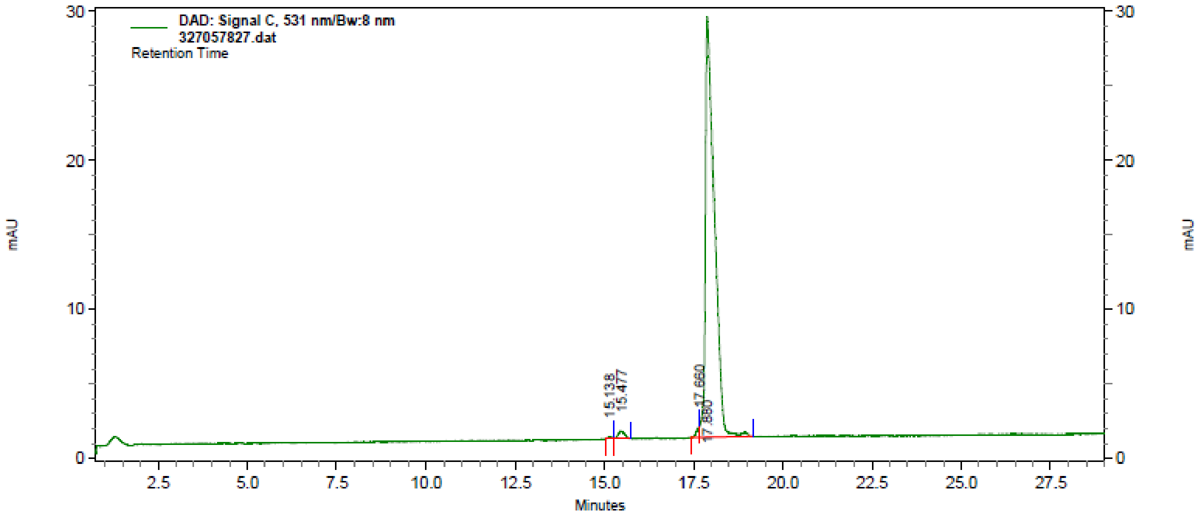


DAD: Signal C, 531
nm/Bw:8 nm Results

| Peak # | Time | Height | Area | Area Percent |
|---|---|---|---|---|
| 1 | 15.14 | 293 | 2387 | 0.2 |
| 2 | 15.48 | 1108 | 14277 | 1.3 |
| 3 | 17.66 | 1460 | 7702 | 0.7 |
| 4 | 17.88 | 59110 | 1033939 | 97.7 |
| Totals | | 61971 | 1058305 | 100.0 |

**Fig. S3. Quality control analysis report of StxSense 2 (integrated DNA Technologies, idt).**

### Ramming et al., Fig. S4

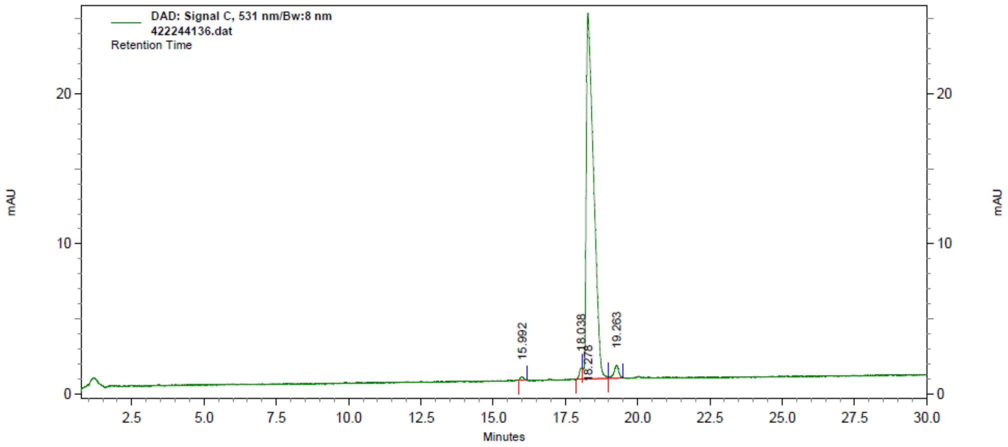


**DAD: Signal C, 531 nm/Bw:8 nm Results**

| Peak # | Time | Height | Area | Area Percent |
|---|---|---|---|---|
| 1 | 15.99 | 483 | 4016 | 0.4 |
| 2 | 18.04 | 1597 | 10996 | 1.2 |
| 3 | 18.28 | 51150 | 900948 | 96.2 |
| 4 | 19.26 | 1827 | 20544 | 2.2 |
| Totals | | 55057 | 936504 | 100.0 |

**Fig. S4. Quality control analysis report of StxSense 3 (integrated DNA Technologies, idt).**

### Ramming et al., Fig. S5

**Integrated DNA Technologies** Page 1 of 1

**Analytical OligoPro CE Report**

Sample ID: 233071385-21 SS HPLC A2 55887
Instrument: Oligo Pro (Offline) Operator: HTA
Acquired: 8/22/2022 5:55:00 AM Reviewed: 8/22/2022 7:48:16 AM

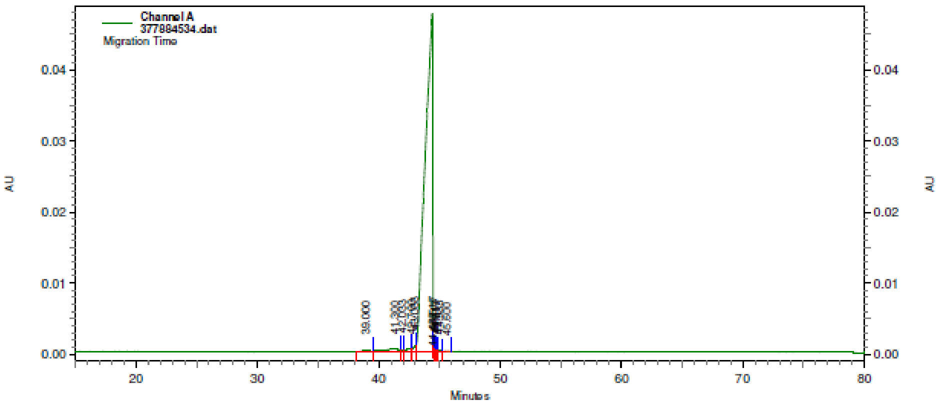


Channel A Results

| Peak # | Time | Height | Corrected Area | Corrected Area Percent |
|---|---|---|---|---|
| 1 | 39.00 | 145 | 160 | 0.4 |
| 2 | 41.30 | 372 | 757 | 1.7 |
| 3 | 42.03 | 284 | 92 | 0.2 |
| 4 | 42.70 | 456 | 294 | 0.7 |
| 5 | 43.03 | 821 | 258 | 0.6 |
| 6 | 44.43 | 47506 | 42884 | 95.9 |
| 7 | 44.52 | 879 | 50 | 0.1 |
| 8 | 44.62 | 433 | 45 | 0.1 |
| 9 | 44.72 | 292 | 43 | 0.1 |
| 10 | 44.98 | 269 | 86 | 0.2 |
| 11 | 45.60 | 70 | 31 | 0.1 |
| Totals | | 51527 | 44700 | 100.0 |

**Fig. S5. Quality control analysis report of StxSense 4 (integrated DNA Technologies, idt).**

### Ramming et al., Fig. S6: SRL substrates (belongs to Fig. 2)

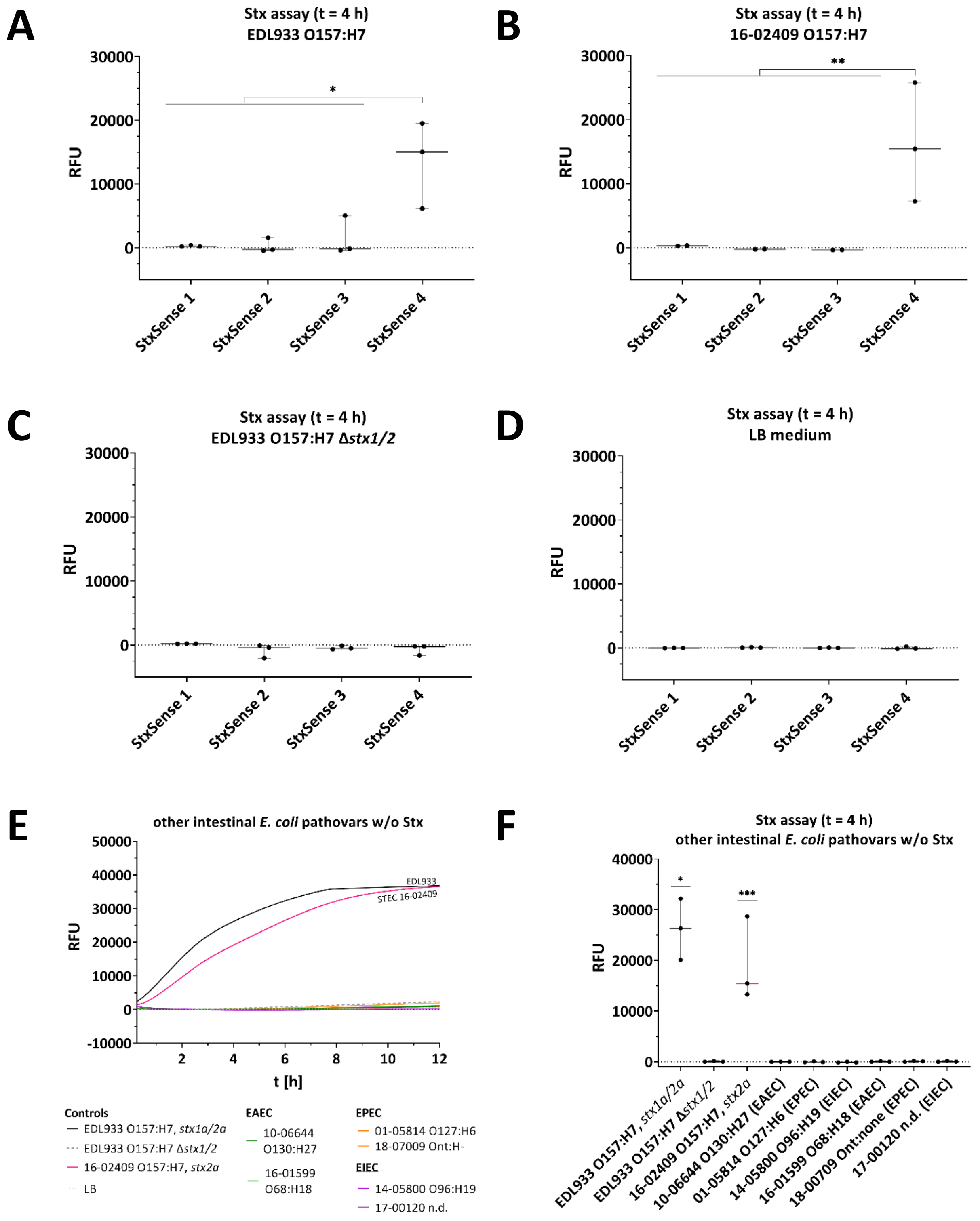

**Ramming et al., Fig. S6 continued**

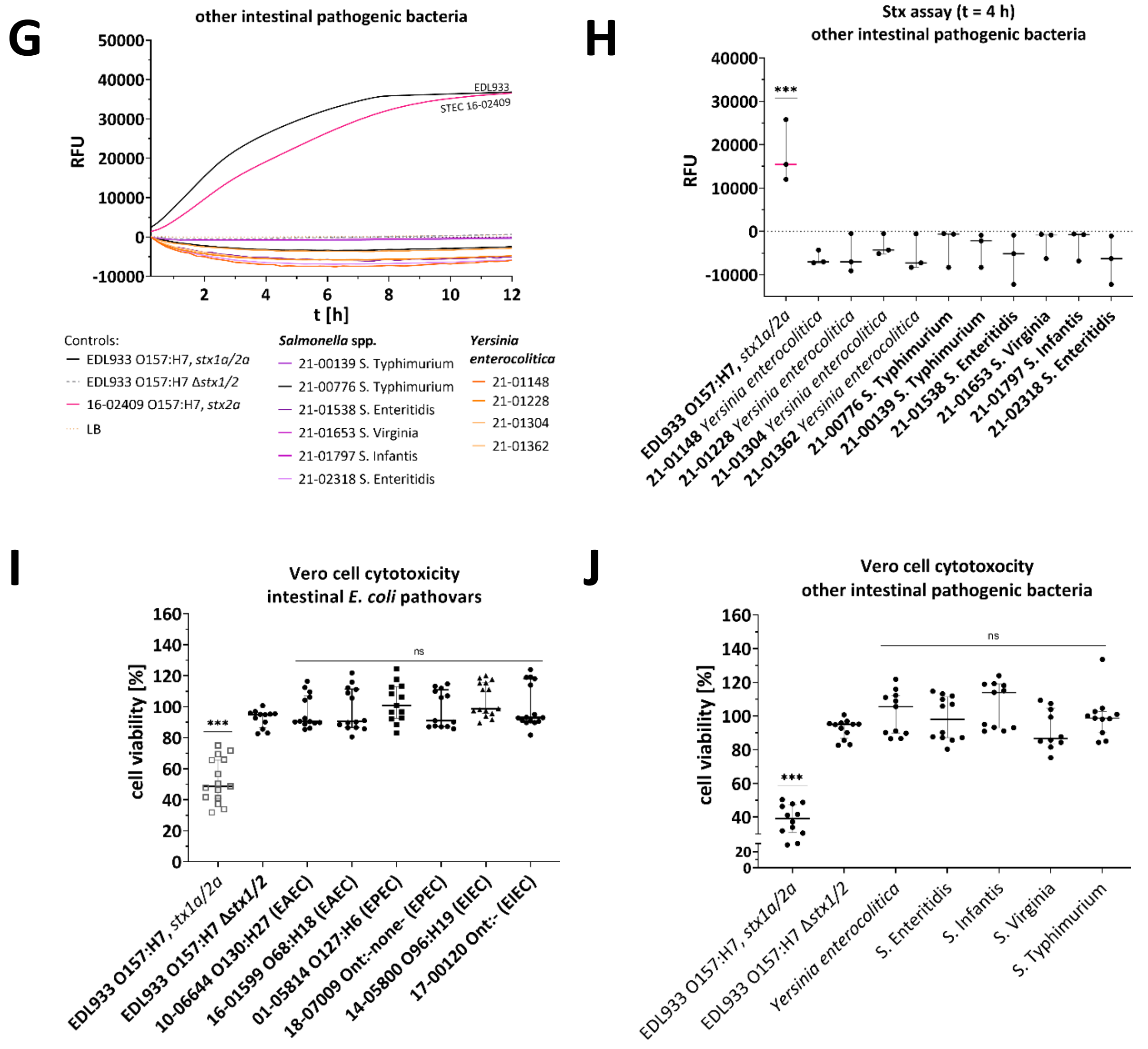


**Fig. S6. Statistics of the Stx enzyme activity assay for STEC and controls. Statistics of the Vero cell cytotoxicity and Stx enzyme activity assay for other intestinal pathogenic bacteria producing no Stx.** Detected fluorescence as a marker of substrate hydrolysis by Stx from different culture supernatants after 4 h incubation of **(A)** EDL933 O157:H7, *stx1a/2a*; **(B)** 16-02409 O157:H7, *stx2a*, **(C)** EDL933 O157:H7 Δ*stx1/2*, and **(D)** LB using the four SRL substrates. Data refer to the data shown in Fig. 2. Other intestinal *E. coli* pathovars: **(E and F)** Stx enzymatic activity assay for culture supernatants for up to 12 h (E) or for 4 h (F) and **(I)** Vero cell cytotoxicity assay. Other intestinal pathogenic bacteria: **(G and H)** Stx enzymatic activity assay for culture supernatants for up to 12 h (H) or for 4 h (I) and **(J)** Vero cell cytotoxicity assay. For Vero cell cytotoxicity assay, cell viability of Vero cells was analyzed using MTT assay after inoculation with diluted bacterial culture supernatants (1:400) for 48 h**.** For statistics of Stx enzymatic activity assay, RFU at 4 h reaction time and 44 °C are shown. The results represent the medians of triplicate samples (n = 3) and are representative of three independent experiments. Error bars represent standard deviation. Statistical analysis was performed by unpaired, double-sided t test (A-D, F, H), Mann-Whitney test (I and J) (*, $p < 0.05$; **, $p < 0.01$; ***, $p < 0.001$), with results compared to those of EDL933 O157:H7 Δ*stx1/2*. RFU, Relative Fluorescent Unit.

### Ramming et al., Fig. S7: assay conditions

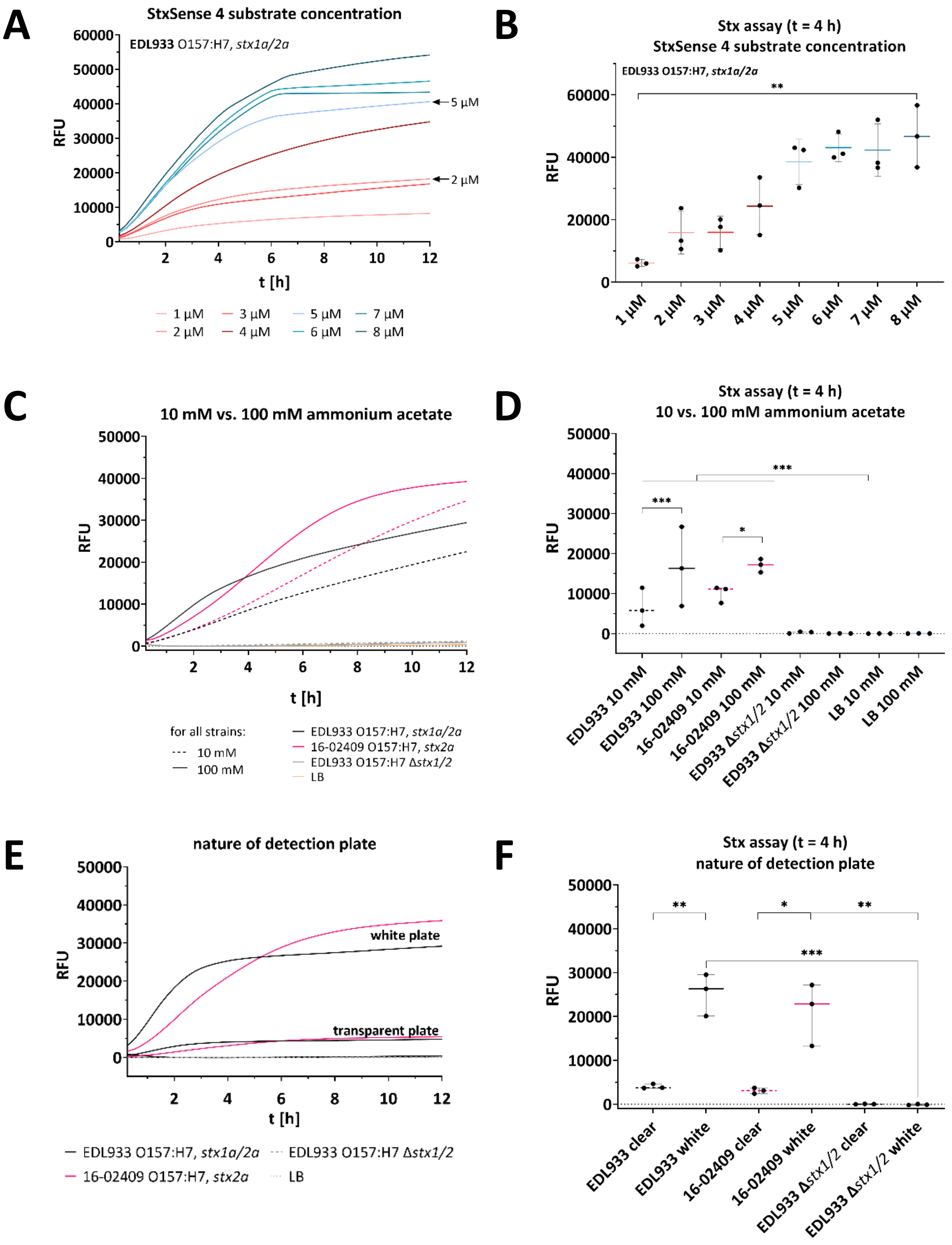

**Ramming et al., Fig. S7 continued**

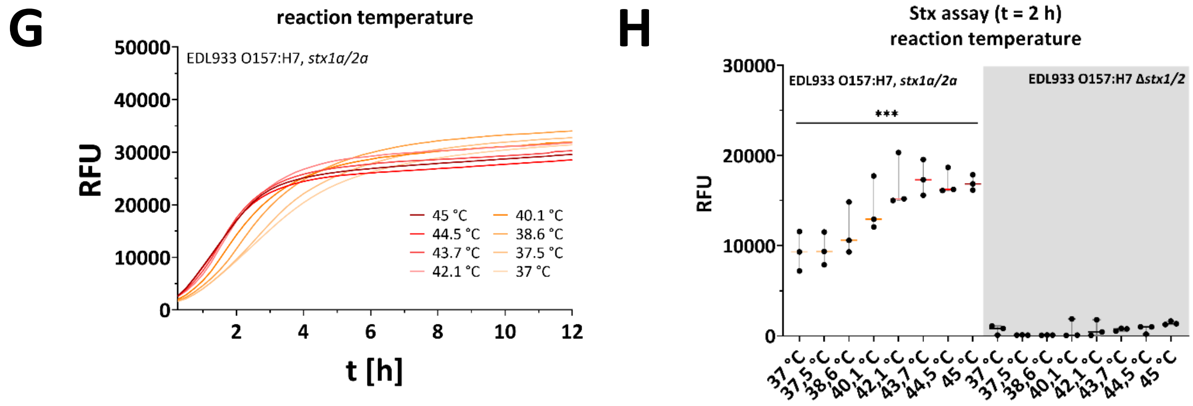


**Fig. S7. Optimal assay conditions for Stx detection in culture supernatants.** Detected fluorescence as a marker of substrate hydrolysis by Stx from positive control EDL933 O157:H7, *stx1a/2a* and/or test strain STEC 16-02409 O157:H7, *stx2a*, and negative control EDL933 O157:H7 Δ*stx1/2*. **(A and B)** Optimal fluorescence readout for Stx was achieved using SRL substrate StxSense 4 concentrations from 2 µM in 100 mM ammonium acetate. **(C and D)** 100 mM ammonium acetate yielded in higher fluorescence signals *stx2a*-producing strain compared to 10 mM ammonium acetate. **(E and F)** Using a white 96 well plate instead of a clear plate was essential for fluorescence detection of Stx-positive samples. **(G and H)** Reaction temperatures above 43.7 °C were optimal Stx detection. Reaction conditions for (A, B, E, F) were 44 °C, 100 mM ammonium acetate, 2 µM StxSense 4. The results represent the medians of triplicate samples (n = 3) and are representative of three independent experiments. Error bars represent standard deviation. Statistical analysis was performed by Mann-Whitney test for non-normally distributed samples (B, H) and unpaired, double-sided t test (D, F) (*, p < 0.05; **, p < 0.01; ***, p < 0.001), with results compared to those of EDL933 O157:H7 Δ*stx1/2* (negative control; -- [grey dashed lines]). RFU, relative fluorescence units; t [h], time [hours].

### Ramming et al., Fig. S8: Stx-producing strains (belongs to Fig. 4)

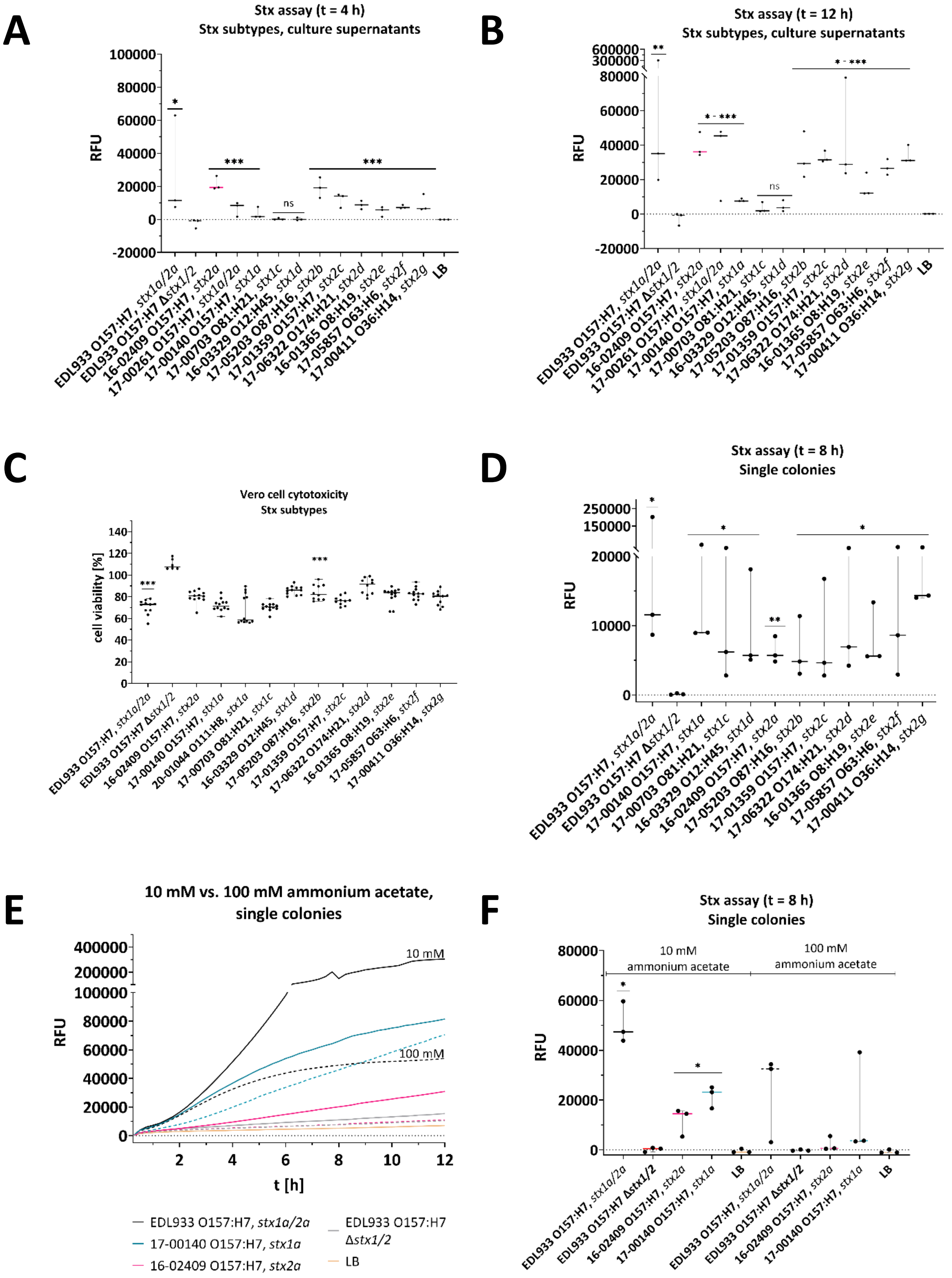

### Ramming et al., Fig. S8 continued

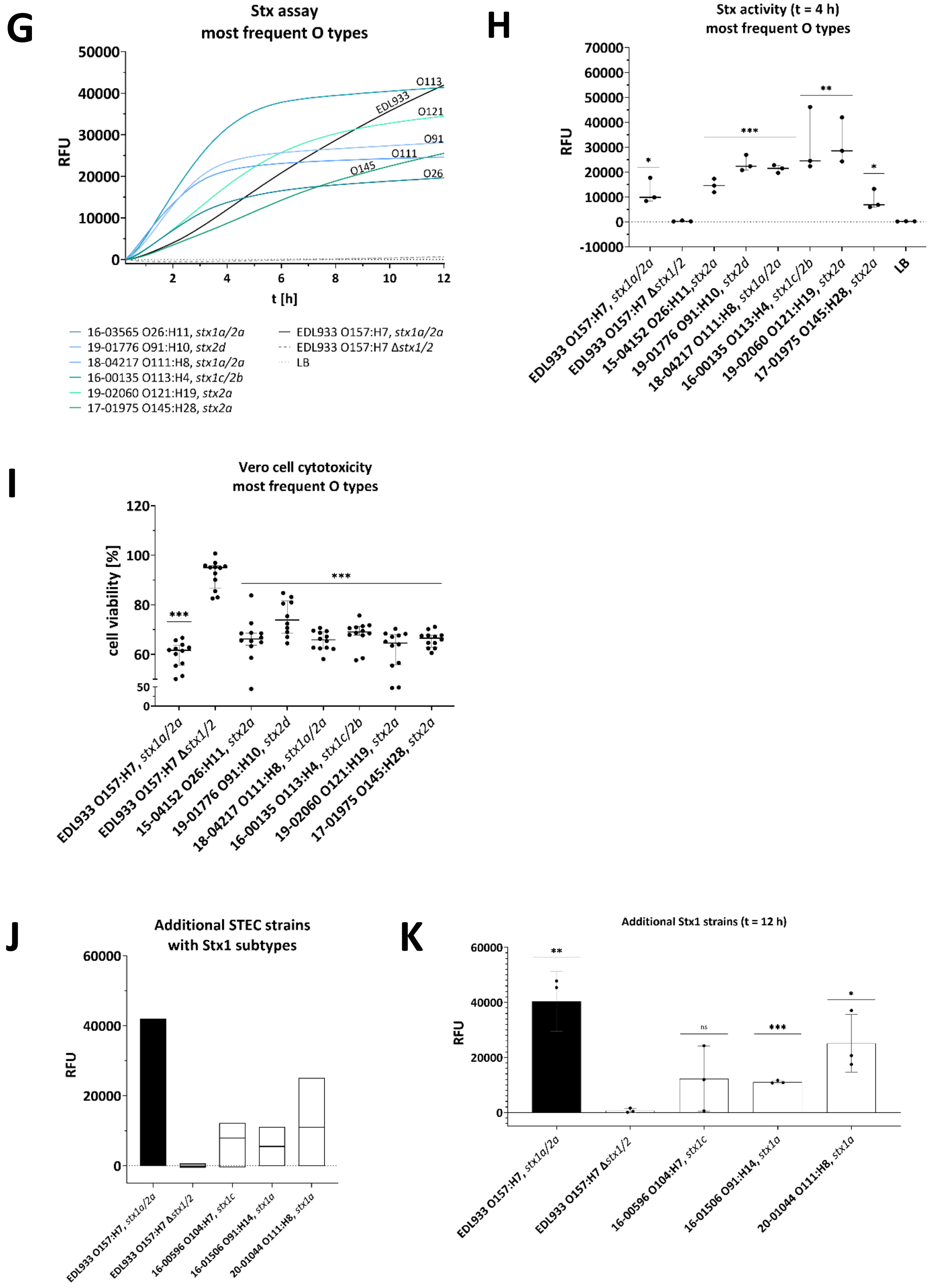

### Ramming et al., Fig. S8 continued (2)

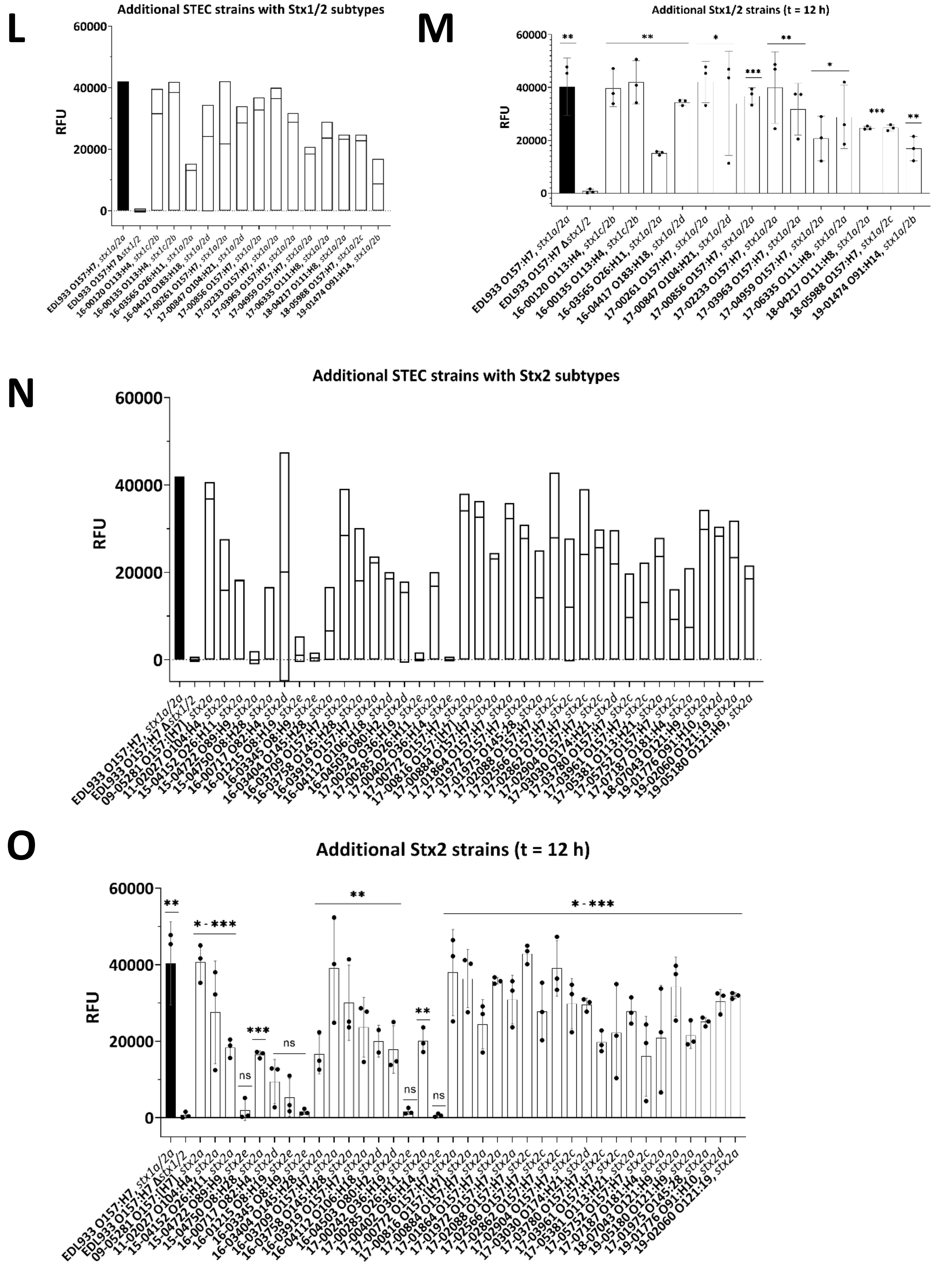

**Ramming et al., Fig. S8 continued (3)**

**P**

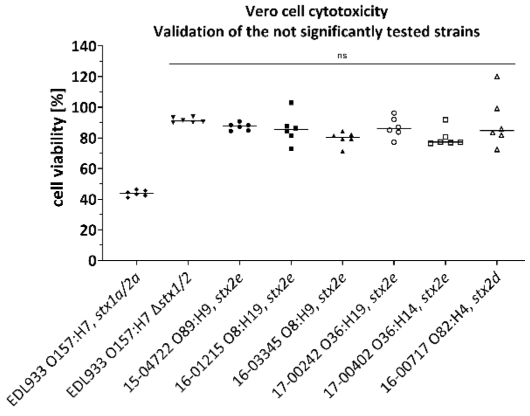


**Fig. S8. Statistics of Vero cell cytotoxicity assay and Stx activity assay for Stx1 and Stx2 subtypes (see Fig. 4A). (A, B, D-H, J-O)** Detected fluorescence of Stx activity from **(A and B)** culture supernatants of STEC strains covering Stx1a-Stx1d and Stx2a-2g at (A) 4 h and (B) 12 h; **(D)** single colonies of STEC strains covering Stx1a-Stx1d and Stx2a-2g, **(E and F)** single colonies of STEC strains and influence of 10 mM vs. 100 mM ammonium acetate, **(G and H)** most frequent serotypes; **(J to O)** Detected fluorescence of culture supernatants of different STEC strains comprising **(J and K)** Stx1, **(L and M)** Stx1/2, **(N and O)** Stx2, with positive control EDL933 O157:H7, *stx1a/2a* (black filled bar), and negative control EDL933 O157:H7 Δ*stx1/2* (grey filled bar) as floating bars (min to max RFU) with line at the median RFU over 12 h and statistics. (J to O) were performed within the same analysis. Shown are the RFU of culture supernatants at 4 h reaction time in 100 mM ammonium acetate, if not stated otherwise, 2 µM StxSense 4, 44 °C, white reaction plate. **(C, I and P)** Cell viability of Vero cells was analyzed using MTT assay after inoculation with diluted bacterial culture supernatants (1:400) for 48 h: **(C)** STEC strains comprising different Stx subtypes, **(I)** most frequent serotypes. Data refer to the data shown in Fig. 4, **(P)** Verification of STEC strains not significantly tested within the STEC detection assay. The results are medians of triplicate samples (n = 3) of three independent experiments. Error bars represent standard deviation. Statistical analysis was performed by Mann-Whitney test (A, B, D-F, H, K, M, O) and unpaired, double-sided t test (C, I) (*, $p < 0.05$; **, $p < 0.01$; ***, $p < 0.001$), with results compared to those of EDL933 O157:H7 Δ*stx1/2*. RFU, Relative Fluorescent Unit; ns, not significant.

**Ramming et al., Fig. S9:** ***Shigella***

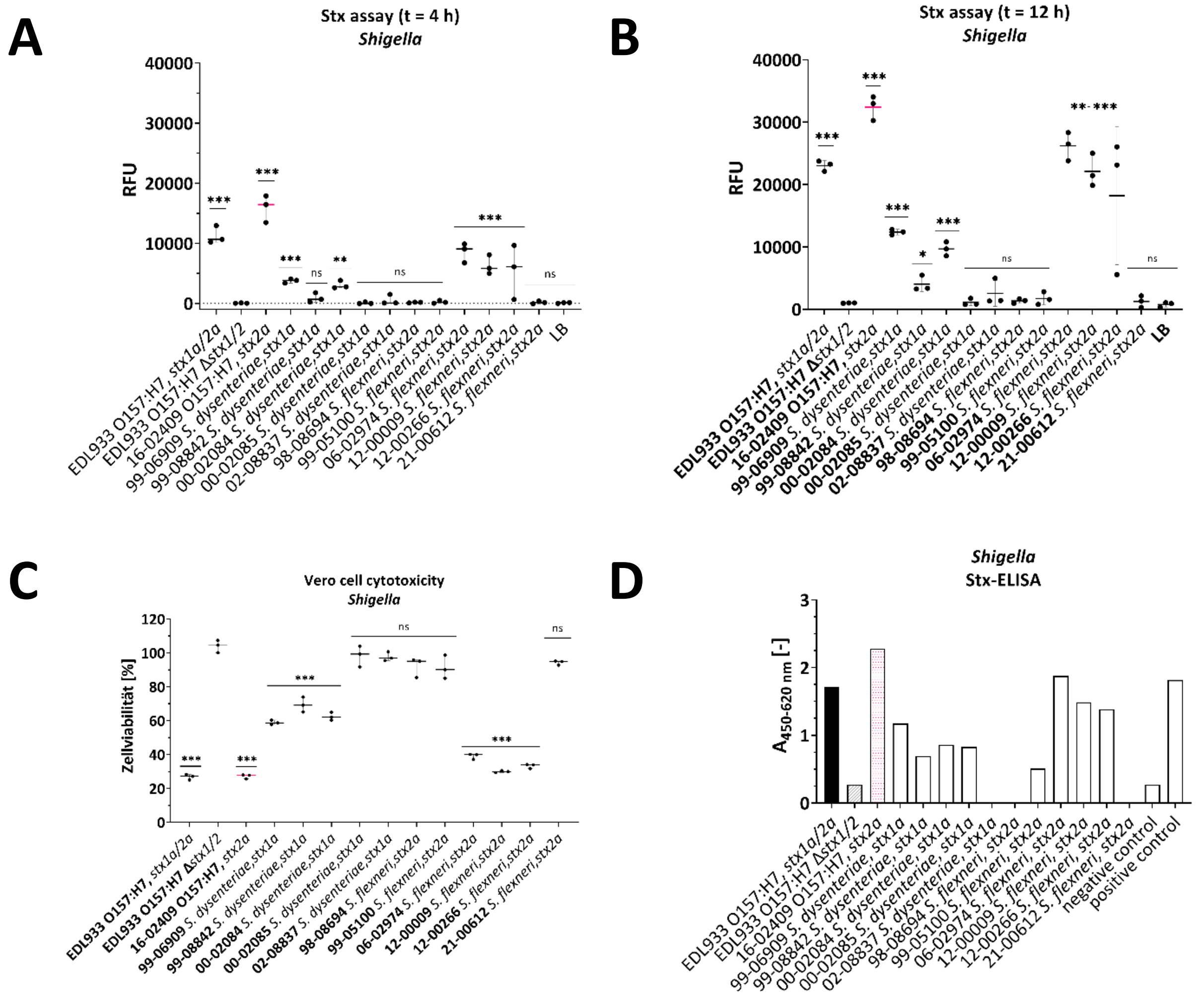


**Fig. S9. Statistics of Vero cell cytotoxicity assay and Stx activity assay for different *Shigella* strains. (A and B)** Detected fluorescence of culture supernatants of different *Shigella* spp. strains and positive control EDL933 O157:H7, *stx1a/2a*, STEC strain 16-02409 O157:H7, *stx2a* and negative control EDL933 O157:H7 Δ*stx1/2* at (A) 4h and (B) 12 h. The assay was performed using 100 mM ammonium acetate, 2 µM StxSense 4, 44 °C. **(C)** Cell viability of Vero cells was analyzed using MTT assay after inoculation with diluted bacterial culture supernatants (1:400) for 48 h. **(D)** Commercial Stx ELISA (R-Biopharm AG, Darmstadt, Germany) of culture supernatant ($OD_{600}$ of 3.0) from different *Shigella* strains detecting Stx1 and Stx2 (n = 1). All other results are medians of triplicate samples (n = 3) of three independent experiments. Error bars represent standard deviation. Statistical analysis was performed by Mann-Whitney test (A) and unpaired, double-sided t test (B) (*, $p < 0.05$; **, $p < 0.01$; ***, $p < 0.001$) with results compared to those of EDL933 O157:H7 Δ*stx1/2*. RFU, Relative Fluorescent Unit; ns, not significant.

**Table S2: STEC strains used in the study (Ramming et al., xxx).**
In total, 65 STEC strains were analyzed for Stx activity. The determined genome sequences were uploaded to the European Nucleotide Archive under the study Acc. No. PRJEB32361 (Lang et al., 2019; Lang et al., 2023). The STEC serotype was determined using Whole Genome Analysis, classical serotyping or PCR-based serotyping (Scheutz et al., 2012; Lang et al., 2019). *eaeA* presence was determined using Whole Genome Analysis or PCR-based serotyping (Schmidt et al., 1993; Lang et al., 2019). n.d., not determined; +, positive analysis test result; -, negative analysis test result; ns, not significant.

| | Strain No. | Year of isolation | O group | H type | MLST sequence type | Accession No. | stx1 subtype | stx2 subtype | *eaeA* | Reference | FRET assay supernatant | where to find? |
|---|---|---|---|---|---|---|---|---|---|---|---|---|
| 1 | **EDL933** | 2001 | 157 | 7 | 11 | GCF_000732965.1 | stx1a | stx2a | + | Perna et al., 2001 | + | pos control |
| 2 | **EDL933 Δ*stx1/2*** | 2007 | 157 | 7 | n.d. | - | - | - | + | Gobert et al., 2007 | - | neg control |
| 3 | **09-05281** | 2009 | 157 | -/[7] | n.d. | n.d. | - | stx2a | + | This study | + | Fig. S8N |
| 4 | **11-02027** | 2011 | 104 | 4 | 678 | SAMN03168461 | - | stx2a | - | Frank et al., 2011 | + | Fig. S8N |
| 5 | **15-04152** | 2015 | 26 | 11 | 29 | ERS3396064 (SAMEA5591849) | - | stx2a | + | Lang et al., 2019 | + | Fig. S8N |
| 6 | **15-04722** | 2015 | 89 | 9 | 10 | ERS3396066 (SAMEA5591851) | - | stx2e | - | Lang et al., 2019 | n.s. | Fig. S8N |
| 7 | **15-04750** | 2015 | 8 | 28 | 162 | ERS3396067 (SAMEA5591852) | - | stx2e | - | Lang et al., 2019 | + | Fig. S8N |
| 8 | **16-00120** | 2016 | 113 | 4 | 10 | ERS3396073 (SAMEA5591858) | stx1c | stx2b | - | Lang et al., 2019 | + | Fig. S8L |
| 9 | **16-00135** | 2016 | 113 | 4 | 10 | ERS3396074 (SAMEA5591859) | stx1c | stx2b | - | Lang et al., 2019 | + | Fig. S8G |
| 10 | **16-00596** | 2016 | 104 | 7 | 1817 | ERS3396083 (SAMEA5591868) | stx1c | - | - | Lang et al., 2019 | + | Fig. S8J |
| 11 | **16-00717** | 2016 | 82 | 4 | 10 | ERS3396090 (SAMEA5591875) | - | stx2d | - | Lang et al., 2019 | n.s. | Fig. S8N |
| 12 | **16-01215** | 2016 | 8 | 19 | 88 | ERS3396098 (SAMEA5591883) | - | stx2e | - | Lang et al., 2019 | n.s. | Fig. S8N |
| 13 | **16-01365** | 2016 | 8 | 19 | 201 | ERS3396100 (SAMEA5591885) | - | stx2e | - | Lang et al., 2019 | + | Fig. 4A/B |
| 14 | **16-01506** | 2016 | 91 | 14 | 33 | ERS3396103 (SAMEA5591888) | stx1a | - | - | Lang et al., 2019 | + | Fig. S8J |
| 15 | **16-02409** | 2016 | 157 | 7 | 11 | ERS3396113 (SAMEA5591898) | - | stx2a | + | Lang et al., 2019 | + | Fig. 2, 4C |
| 16 | **16-03329** | 2016 | 12 | 45 | n.d. | ERS3396123 (SAMEA5591908) | stx1d | - | - | Lang et al., 2019 | + | Fig. 4A/B |
| 17 | **16-03345** | 2016 | 8 | 9 | 23 | ERS3396125 (SAMEA5591910) | - | stx2e | - | Lang et al., 2019 | n.s. | Fig. S8N |
| 18 | **16-03404** | 2016 | 145 | 28 | 32 | ERS3396126 (SAMEA5591911) | - | stx2a | + | Lang et al., 2019 | + | Fig. S8N |
| 19 | **16-03565** | 2016 | 26 | 11 | 21 | ERS3396129 (SAMEA5591914) | stx1a | stx2a | + | Lang et al., 2019 | + | Fig. S8G |
| 20 | **16-03709** | 2016 | 157 | 7 | 11 | ERS3396134 (SAMEA5591919) | - | stx2a | + | Lang et al., 2019 | + | Fig. S8N |
| 21 | **16-03758** | 2016 | 145 | 28 | 32 | ERS3396135 (SAMEA5591920) | - | stx2a | + | Lang et al., 2019 | + | Fig. S8N |
| 22 | **16-03919** | 2016 | 157 | 7 | 11 | ERS3396136 (SAMEA5591921) | - | stx2a | + | Lang et al., 2019 | + | Fig. S8N |
| 23 | **16-04112** | 2016 | 106 | 18 | 663 | ERS3396140 (SAMEA5591925) | - | stx2d | - | Lang et al., 2019 | + | Fig. S8N |
| 24 | **16-04417** | 2016 | 183 | 18 | 657 | ERS3396151 (SAMEA5591936) | stx1a | stx2d | - | Lang et al., 2019 | + | Fig. S8L |
| 25 | **16-04503** | 2016 | 80 | 2 | 301 | ERS3396152 (SAMEA5591937) | - | stx2d | + | Lang et al., 2019 | + | Fig. S8N |
| 26 | **17-00140-1** | 2017 | 157 | 7 | 11 | ERS3396169 (SAMEA5591954) | stx1a | - | + | Lang et al., 2019 | + | Fig. 4A/B |
| 27 | **17-00242** | 2017 | 36 | 19 | 10 | ERS3396170 (SAMEA5591955) | - | stx2e | - | Lang et al., 2019 | n.s. | Fig. S8L |
| 28 | **17-00261** | 2017 | 157 | 7 | 11 | ERS3396171 (SAMEA5591956) | stx1a | stx2a | + | Lang et al., 2019 | + | Fig. 4A/B |
| 29 | **17-00285** | 2017 | 26 | 11 | 29 | ERS3396172 (SAMEA5591957) | - | stx2a | + | Lang et al., 2019 | + | Fig. S8L |
| 30 | **17-00402** | 2017 | 36 | 14 | 1176 | ERS3396173 (SAMEA5591958) | - | stx2e | - | Lang et al., 2019 | n.s. | Fig. S8L |
| 31 | **17-00411** | 2017 | 36 | 14 | 1176 | ERS3396174 (SAMEA5591959) | - | stx2g | - | Lang et al., 2019 | + | Fig. 4A/B |
| 32 | **17-00703** | 2017 | 81 | 21 | 737 | ERS3396178 (SAMEA5591963) | stx1c | - | - | Lang et al., 2019 | + | Fig. 4A/B |
| 33 | **17-00772** | 2017 | 157 | 7 | 11 | ERS3396179 (SAMEA5591964) | - | stx2a | + | Lang et al., 2019 | + | Fig. S8N |
| 34 | **17-00816** | 2017 | 157 | -/[7] | n.d. | n.d. | - | stx2a | + | This study | + | Fig. S8N |
| 35 | **17-00847** | 2017 | 104 | 21 | 672 | ERS3396182 (SAMEA5591967) | stx1a | stx2d | - | Lang et al., 2019 | + | Fig. S8L |
| 36 | **17-00856** | 2017 | 157 | 7 | 11 | ERS3396183 (SAMEA5591968) | stx1a | stx2a | + | Lang et al., 2019 | + | Fig. S8L |
| 37 | **17-00884** | 2017 | 157 | 7 | 11 | ERS3396185 (SAMEA5591970) | - | stx2a | + | Lang et al., 2019 | + | Fig. S8N |
| 38 | **17-01359** | 2017 | 157 | 7 | 11 | ERS3396196 (SAMEA5591981) | - | stx2c | + | Lang et al., 2019 | + | Fig. 4A/B |
| 39 | **17-01864** | 2017 | 157 | 7 | 11 | ERS3396199 (SAMEA5591984) | - | stx2a | + | Lang et al., 2019 | + | Fig. S8N |
| 40 | **17-01972** | 2017 | 157 | 7 | 587 | ERS3396200 (SAMEA5591985) | - | stx2a | + | Lang et al., 2019 | + | Fig. S8N |
| 41 | **17-01975** | 2017 | 145 | 28 | 32 | ERS3396201 (SAMEA5591986) | - | stx2a | + | Lang et al., 2019 | + | Fig. S8G |
| 42 | **17-02088** | 2017 | 157 | 7 | 1804 | ERS3396202 (SAMEA5591987) | - | stx2c | + | Lang et al., 2019 | + | Fig. S8N |
| 43 | **17-02233** | 2017 | 157 | 7 | 11 | ERS3396203 (SAMEA5591988) | stx1a | stx2a | + | Lang et al., 2019 | + | Fig. S8L |
| 44 | **17-02566** | 2017 | 157 | 7 | 11 | ERS3396204 (SAMEA5591989) | - | stx2c | + | Lang et al., 2019 | + | Fig. S8N |
| 45 | **17-02862** | 2017 | 157 | 7 | 11 | ERS3396210 (SAMEA5591995) | - | stx2c | + | Lang et al., 2019 | + | Fig. S8N |
| 46 | **17-02904** | 2017 | 157 | 7 | n.d. | n.d. | - | stx2c | + | This study | + | Fig. S8N |
| 47 | **17-03030** | 2017 | 174 | 21 | 677 | ERS3396212 (SAMEA5591997) | - | stx2d | - | Lang et al., 2019 | + | Fig. S8N |
| 48 | **17-03780** | 2017 | 157 | 7 | 11 | ERS3396215 (SAMEA5592000) | - | stx2c | + | Lang et al., 2019 | + | Fig. S8N |
| 49 | **17-03961** | 2017 | 157 | 7 | 11 | ERS3396218 (SAMEA5592003) | - | stx2c | + | Lang et al., 2019 | + | Fig. S8N |
| 50 | **17-03963** | 2017 | 157 | 7 | 11 | ERS3396219 (SAMEA5592004) | stx1a | stx2a | + | Lang et al., 2019 | + | Fig. S8L |
| 51 | **17-04959** | 2017 | 157 | 7 | 11 | ERS3396225 (SAMEA5592010) | stx1a | stx2a | + | Lang et al., 2019 | + | Fig. S8L |
| 52 | **17-05203** | 2017 | 87 | 16 | 2101 | ERS3396229 (SAMEA5592014) | - | stx2b | - | Lang et al., 2019 | + | Fig. 4A/B |
| 53 | **17-05381** | 2017 | 113 | 21 | 223 | ERS3396236 (SAMEA5592021) | - | stx2a | - | Lang et al., 2019 | + | Fig. S8N |
| 54 | **17-05752** | 2017 | 157 | 7 | 11 | ERS3396247 (SAMEA5592032) | - | stx2c | + | Lang et al., 2019 | + | Fig. S8N |
| 55 | **17-05857** | 2017 | 63 | 6 | 583 | ERS3396253 (SAMEA5592038) | - | stx2f | + | Lang et al., 2019 | + | Fig. 4A/B |
| 56 | **17-06322** | 2017 | 174 | 21 | 677 | ERS3396266 (SAMEA5592051) | - | stx2d | - | Lang et al., 2019 | + | Fig. 4A/B |
| 57 | **17-06335** | 2017 | 111 | 8 | 16 | ERS3396269 (SAMEA5592054) | stx1a | stx2a | + | Lang et al., 2019 | + | Fig. S8L |
| 58 | **17-07187** | 2017 | 181 | 4 | n.d. | n.d. | - | stx2a | - | Lang et al., 2023 | + | Fig. S8N |
| 59 | **18-04217** | 2018 | 111 | 8 | n.d. | n.d. | stx1a | stx2a | + | This study | + | Fig. S8G |
| 60 | **18-05988** | 2018 | 157 | 7 | n.d. | n.d. | stx1a | stx2c | + | This study | + | Fig. S8L |
| 61 | **18-07043** | 2018 | 121 | 9 | n.d. | n.d. | - | stx2a | + | This study | + | Fig. S8N |
| 62 | **19-01474** | 2019 | 91 | 14 | n.d. | n.d. | stx1a | stx2b | - | This study | + | Fig. S8L |
| 63 | **19-01776** | 2019 | 91 | 10 | n.d. | n.d. | - | stx2d | - | This study | + | Fig. S8G |
| 64 | **19-02060** | 2019 | 121 | 19 | n.d. | n.d. | - | stx2a | + | This study | + | Fig. S8G |
| 65 | **19-05180** | 2019 | 121 | 19 | n.d. | n.d. | - | stx2a | + | This study | + | Fig. S8N |
| 66 | **20-01044** | 2020 | 111 | 8 | n.d. | n.d. | stx1a | - | + | This study | + | Fig. S8J |

**Table S3: Other intestinal *E. coli* pathovar strains without Stx used in the study (Ramming et al., xxx).**

The *E. coli* pathovar serotype was determined using Whole Genome Analysis, classical serotyping or PCR-based serotyping (Scheutz et al., 2012; Lang et al., 2019). *eaeA* presence was determined using Whole Genome Analysis or PCR-based serotyping (Schmidt et al., 1993; Lang et al., 2019). n.d., not determined; +, positive analysis test result; -, negative analysis test result.

| | **Strain no.** | **Year of isolation** | **Pheno-typic O group** | **Pheno-typic H type** | **pathovar** | ***eaeA*** | **Reference** | **FRET assay supernatant** | **where to find?** | **Vero cell cytotoxicity assay** | **where to find?** |
|---|---|---|---|---|---|---|---|---|---|---|---|
| 1 | **10-06644** | 2010 | 130 | 27 | EAEC | n.d. | This study | - | Fig. S6E, F | - | Fig. S6I |
| 2 | **16-01599-2** | 2016 | 68 | 18 | EAEC | - | This study | - | Fig. S6E, F | - | Fig. S6I |
| 3 | **01-05814** | 2001 | 127 | 6 | EPEC | + | This study | - | Fig. S6E, F | - | Fig. S6I |
| 4 | **18-07009-2** | 2018 | Ont | - | EPEC | + | This study | - | Fig. S6E, F | - | Fig. S6I |
| 5 | **18-07383-2** | 2018 | Ont | - | EPEC | + | This study | - | Fig. S6E, F | - | Fig. S6I |
| 6 | **14-05800-2** | 2014 | 96 | 19 | EIEC | n.d. | This study | - | Fig. S6E, F | - | Fig. S6I |
| 7 | **17-00120** | 2017 | n.d. | n.d. | EIEC | n.d. | This study | - | Fig. S6E, F | - | Fig. S6I |

**Table S4: *Yersinia* and *Salmonella* strains used in the study (Ramming et al., xxx).**

The species or serovar was determined using Whole Genome Analysis, classical serotyping or PCR-based serotyping (Lang et al., 2019). +, positive analysis test result; -, negative analysis test result.

| | Strain no. | Year of isolation | Species or serovar | Stx1 subtype | Stx2 subtype | Reference | FRET assay supernatant | where to find? | Vero cell cytotoxicity assay | where to find? |
|---|---|---|---|---|---|---|---|---|---|---|
| 1 | **21-01148** | 2021 | *Yersinia enterocolitica* | - | - | This study | - | Fig. S6G, H | - | Fig. S6J |
| 2 | **21-01228** | 2021 | *Yersinia enterocolitica* | - | - | This study | - | Fig. S6G, H | - | Fig. S6J |
| 3 | **21-01304** | 2021 | *Yersinia enterocolitica* | - | - | This study | - | Fig. S6G, H | - | Fig. S6J |
| 4 | **21-01362** | 2021 | *Yersinia enterocolitica* | - | - | This study | - | Fig. S6G, H | - | Fig. S6J |
| 5 | **21-00139** | 2021 | *Salmonella enterica* S. Typhimurium | - | - | This study | - | Fig. S6G, H | - | Fig. S6J |
| 6 | **21-00776** | 2021 | *Salmonella enterica* S. Typhimurium | - | - | This study | - | Fig. S6G, H | - | Fig. S6J |
| 7 | **21-01538** | 2021 | *Salmonella enterica* S. Enteritidis | - | - | This study | - | Fig. S6G, H | - | Fig. S6J |
| 8 | **21-01653** | 2021 | *Salmonella enterica* S. Virginia | - | - | This study | - | Fig. S6G, H | - | Fig. S6J |
| 9 | **21-01797** | 2021 | *Salmonella enterica* S. Infantis | - | - | This study | - | Fig. S6G, H | - | Fig. S6J |
| 10 | **21-02318** | 2021 | *Salmonella enterica* S. Enteritidis | - | - | This study | - | Fig. S6G, H | - | Fig. S6J |

**Table S4:** ***Shigella*** **strains used in the study (Ramming et al., xxx).**

The species or serovar was determined using Whole Genome Analysis, classical serotyping or PCR-based serotyping (Lang et al., 2019). +, positive analysis test result; -, negative analysis test result.

| | Strain no. | Year of isolation | Species or serovar | Stx1 subtype | Stx2 subtype | Reference | FRET assay supernatant | where to find? | Vero cell cytotoxicity assay | where to find? | Stx ELISA | where to find? |
|---|---|---|---|---|---|---|---|---|---|---|---|---|
| 1 | **99-06909** | 1999 | *Shigella dysenteriae* | stx1a | - | This study | + | Fig. S9A, B | + | Fig. S9C | + | Fig. S9D |
| 2 | **99-08842** | 1999 | *Shigella dysenteriae* | stx1a | - | This study | + | Fig. S9A, B | + | Fig. S9C | + | Fig. S9D |
| 3 | **00-02084** | 2000 | *Shigella dysenteriae* | stx1a | - | This study | + | Fig. S9A, B | + | Fig. S9C | + | Fig. S9D |
| 4 | **00-02085** | 2000 | *Shigella dysenteriae* | stx1a | - | This study | - | Fig. S9A, B | - | Fig. S9C | + | Fig. S9D |
| 5 | **02-08837** | 2002 | *Shigella dysenteriae* | stx1a | - | This study | - | Fig. S9A, B | - | Fig. S9C | - | Fig. S9D |
| 6 | **98-08694** | 1998 | *Shigella flexneri* | - | stx2a | This study | - | Fig. S9A, B | - | Fig. S9C | - | Fig. S9D |
| 7 | **99-05100** | 1999 | *Shigella flexneri* | - | stx2a | This study | - | Fig. S9A, B | - | Fig. S9C | + | Fig. S9D |
| 8 | **06-02974** | 2006 | *Shigella flexneri* | - | stx2a | This study | + | Fig. S9A, B | + | Fig. S9C | + | Fig. S9D |
| 9 | **12-00009** | 2012 | *Shigella flexneri* | - | stx2a | This study | + | Fig. S9A, B | + | Fig. S9C | + | Fig. S9D |
| 10 | **12-00266** | 2012 | *Shigella flexneri* | - | stx2a | This study | + | Fig. S9A, B | + | Fig. S9C | + | Fig. S9D |
| 11 | **21-00612** | 2021 | *Shigella flexneri* | - | stx2a | This study | - | Fig. S9A, B | - | Fig. S9C | - | Fig. S9D |

**Supplemental Methods**

**Western blot analysis**. STEC culture supernatants or cell lysates were electrophoretically separated on a 15 % SDS-polyacrylamide gel. Proteins were transferred to a methanol (99.9 %, Carl Roth GmbH, Karlsruhe, Germany)-activated nitrocellulose membrane (Merck Millipore; Darmstadt, Germany). The membrane was then blocked in 5 % milk (in TBS-T, Sigma-Aldrich, Darmstadt, Germany) at 4 °C overnight. Primary monoclonal mouse antibodies against Stx1 (MBS5307327; MyBioSource Inc., San Diego, USA) or Stx2 (VT 135/6-B9; Sifin, Berlin, Germany) were added at a dilution of 1:1,000 for 3 h. StarBright Blue 520 Conjugated Goat Anti-Mouse Fluorescent Secondary Antibodies (Bio-Rad Laboratories; Feldkirchen, Germany) were used as secondary antibody at a dilution of 1:50,000 for 1 h. Stx was detected by fluorescence at 520 nm (ChemiDoc MP, Bio-Rad Laboratories, StarBright 520 channel).

**Vero cell cytotoxicity assay.** Toxicity of Stx towards Vero cells was determined as described previously (Roberts et al., 2001) with modifications. Prior to addition of culture supernatants, $1x10^5$ Vero cells/mL were seeded into a 96 well-plate (Greiner Bio-One; Kremsmünster, Austria) and were incubated for 24h at 37 °C and 5 % $CO_2$. Culture supernatants were diluted 1:400 in DMEM with 10 % FBS and 200 µL were added to each well and incubated for 48 h at 37 °C and under 5 % $CO_2$. After incubation, the cells were washed with 1× PBS and 100 µL 1-(4,5-Dimethylthiazol-2-yl)-3,5-diphenylformazan (MTT; 0.5 mg/mL in PBS; Sigma) were added for 1 h. After removal, 100 µL acidified isopropanol (isopropanol with 4 % v/v HCl, 32 %) were added. The absorption at 570 nm was photometrically measured at 570 nm (Tecan Infinite M-1000 Pro; Tecan, Suisse) (Mosmann, 1983). The proportion of damaged to viable cells (viability, in %) was calculated using mean absorbance of sample/mean absorbance of Vero cell control × 100.
